# Elevated recessive lethal frequencies drive hatching failure following near extinction in ‘Alalā, the Hawaiian crow

**DOI:** 10.64898/2026.03.24.713968

**Authors:** Christopher C. Kyriazis, Stefanie Grosser, Yasmin Foster, Bryce Masuda, Alison M. Flanagan, Jennifer Balacco, Emma B. Crago, Erin Datlof, Olivier Fedrigo, Giulio Formenti, Catherine E. Grueber, Jacqueline A. Robinson, Jolene T. Sutton, Alan Tracey, Jonathan M.D. Wood, Erich D. Jarvis, Oliver A. Ryder, Bruce C. Robertson, Aryn P. Wilder

## Abstract

Near-extinction events impose severe genomic bottlenecks that can have lasting fitness consequences, yet the specific mechanisms involved remain poorly understood. ‘Alalā, the last remaining Hawaiian crow, narrowly avoided extinction when a conservation breeding program was initiated from just eight genetic founders. Although the conservation breeding program has since recovered to ∼120 birds, it remains plagued by egg hatching failure rates >50%. To investigate the impacts of this bottleneck on hatching failure and other fitness components, we generated a chromosome-level reference genome and resequenced 175 individuals, including 78 deceased embryos. Although long runs of homozygosity (ROH) >1Mb are abundant in ‘Alalā (mean F_ROH_=0.32), associations between F_ROH_ and measures of survival and reproduction, including egg failure, were weak or nonexistent. Instead, we identify two recessive lethal haplotypes that together account for nearly all of the excess hatching failure and have persisted in the population at high frequency (15-25%), hinting at impaired purifying selection. Eco-evolutionary simulation models demonstrate that these findings are expected for a species that has endured a severe population bottleneck and exhibits modest levels of pre-bottleneck heterozygosity. Our findings suggest that elevated recessive allele frequencies may be a broadly important consequence of population bottlenecks.

## Introduction

Mounting anthropogenic pressures have resulted in a growing number of species that are nearing the brink of extinction ^1^. Although targeted conservation efforts are frequently successful at recovering critically endangered species ^2,3^, severe population bottlenecks can impart a genomic cost that may impair recovery and have long-lasting impacts on fitness and genetic diversity ^4,5^. Thus, detailing the genomic mechanisms underlying fitness declines from population bottlenecks is a critical step toward ensuring a robust recovery. This understanding is particularly pressing given that conservation breeding programs require enormous and sustained investment in specialized facilities and personnel, making it all the more urgent to proactively understand the genomic factors that may limit their success.

In birds, numerous species have recovered from severe population bottlenecks, including the Chatham Island black robin (*Petroica traversi*; recovering from just 2 individuals), Mauritius kestrel (*Falco punctatus*; 4 individuals), and California condor (*Gymnogyps californianus*; 22 individuals) ^3^. A growing literature of genomic studies has highlighted the effects of these bottlenecks in driving increased homozygosity and potentially compromising recovery ^6–13^. An important element mediating the fitness impacts of such bottlenecks is the load of segregating (partially) recessive deleterious variation in a species (i.e., inbreeding load or masked load)^14,15^, which is primarily determined by historical demography and mutation rates. For instance, in the Japanese wood pigeon, recovery was enabled in part by the naturally small historical population size, resulting in an absence of detectable inbreeding depression ^11^.

Understanding whether and how species can recover from severe population declines is a particularly pressing issue in Hawai‘i, where ∼75% of native birds are already extinct, and most remaining native species are highly endangered ^16,17^. ‘Alalā, also known as Hawaiian crow (*Corvus hawaiiensis*), was considered abundant across Hawai‘i Island until the end of the 19th century, but declined precipitously during the 20th century and was ultimately declared extinct in the wild in 2002 ^18,19^. Although conservation breeding management has since recovered the population to ∼120 individuals from eight genetic founders, this severe population bottleneck resulted in substantial loss of genetic diversity ^18^ and potentially hindered survival in the conservation breeding program ^20,21^. In particular, ‘Alalā suffer from a low hatching success rate, with >50% of embryos dying before hatching, a problem that has persisted since the start of intensive conservation breeding thirty years ago and continues to limit the reproductive capacity of the ex situ population ^21^

Elevated hatching failure is a well-documented issue for endangered and captive bird species^22,23^. A recent meta-analysis estimated mean hatching failure rates of ∼38% for captive birds, driven largely by non-genetic factors including artificial incubation and limited mate choice ^23^. The high rate of egg hatching failure in ‘Alalā (>50%) notably exceeds this baseline for captive birds, and previous studies have suggested that inbreeding may be a contributing factor ^20,21^. Here, we investigate the genomic basis of hatching failure and other fitness-related traits in ’Alalā, with the goal of informing management strategies that could improve reproductive success and ultimately support the recovery of a wild population. Our findings carry implications not only for ’Alalā conservation but for the broader understanding of how severe population bottlenecks shape the genomic architecture of fitness in endangered species.

## Results

### Reference genome assembly and annotation

To enable a population genomic analysis of ‘Alalā, we first assembled a chromosome-level reference genome using the Vertebrate Genomes Project (VGP) assembly pipeline v2.0, using a combination of PacBio HiFi, Hi-C, and bionano data^24^, followed by manual curation. The resulting assembly (bCorHaw1.pri.cur) was 1.2 Gb in length, consisting of 43 chromosomes and 187 scaffolds with a scaffold N50 of 76.3 Mb, a substantial improvement over the previous assembly that had a contig N50 of 7.7 Mb though was not scaffolded ^25^. Annotation complemented with RNAseq data revealed 21,647 genes, of which 16,414 were protein-coding genes. We determined that 99.1% of avian single-copy gene orthologs were present and complete using BUSCO ^26^. We observed a high degree of synteny between ‘Alalā and three other corvid species (several sequenced also using the VGP pipeline), though with the notable exception of chromosome 2 from other corvid species being split into three chromosomes in ‘Alalā (**Fig. S1**). Similar chromosomal fissions of ancestral macrochromosomes have been documented in other passerine lineages ^27^, suggesting these rearrangements arise recurrently during avian genome evolution. Our genome has become the public reference for the species in NCBI and is contributed to the VGP Phase 1 project of assemblies ^28^.

### High inbreeding and low genome-wide diversity in ‘Alalā

We generated a high-coverage (average depth after filtering = 18.3x) resequencing dataset of genomes of 175 individuals in the conservation breeding program between 1989-2019, including 78 embryos that failed to hatch and 97 hatched individuals that survived for 3 months to 29 years (**Table S1**). Across all individuals in our dataset, we observe a high abundance of runs of homozygosity (ROH), with a mean inbreeding coefficient (F_ROH_) for ROH >1 Mb of 0.32 (range 0.15-0.49; **Figs. 1 & S2**). Levels of inbreeding increased over time in the captive population and were substantially higher than pedigree-based measures of F (F_ped_; **Figs. 2A-B**). Notably, even individuals from the first several generations of the breeding program, when F_ped_ was often quite low, exhibited elevated F_ROH_, suggesting that ROH likely accumulated in the wild ‘Alalā population during a prolonged decline. When lowering the ROH threshold to 100kb, F_ROH_ increases to 0.40 on average (**Fig. S3**), indicative of autozygosity due to longer-term population declines. Finally, we observe that a large proportion of ROH in ‘Alalā are very long (>10 Mb), with an average F_ROH_ for ROH >10 Mb of 0.14, which was more similar in magnitude to F_ped_ (**Fig. S4**), as expected. In many cases, ROH exceeded 50 Mb in length and encompassed nearly the entire chromosome (**Fig. 1 & S2**).

**Figure 1:**
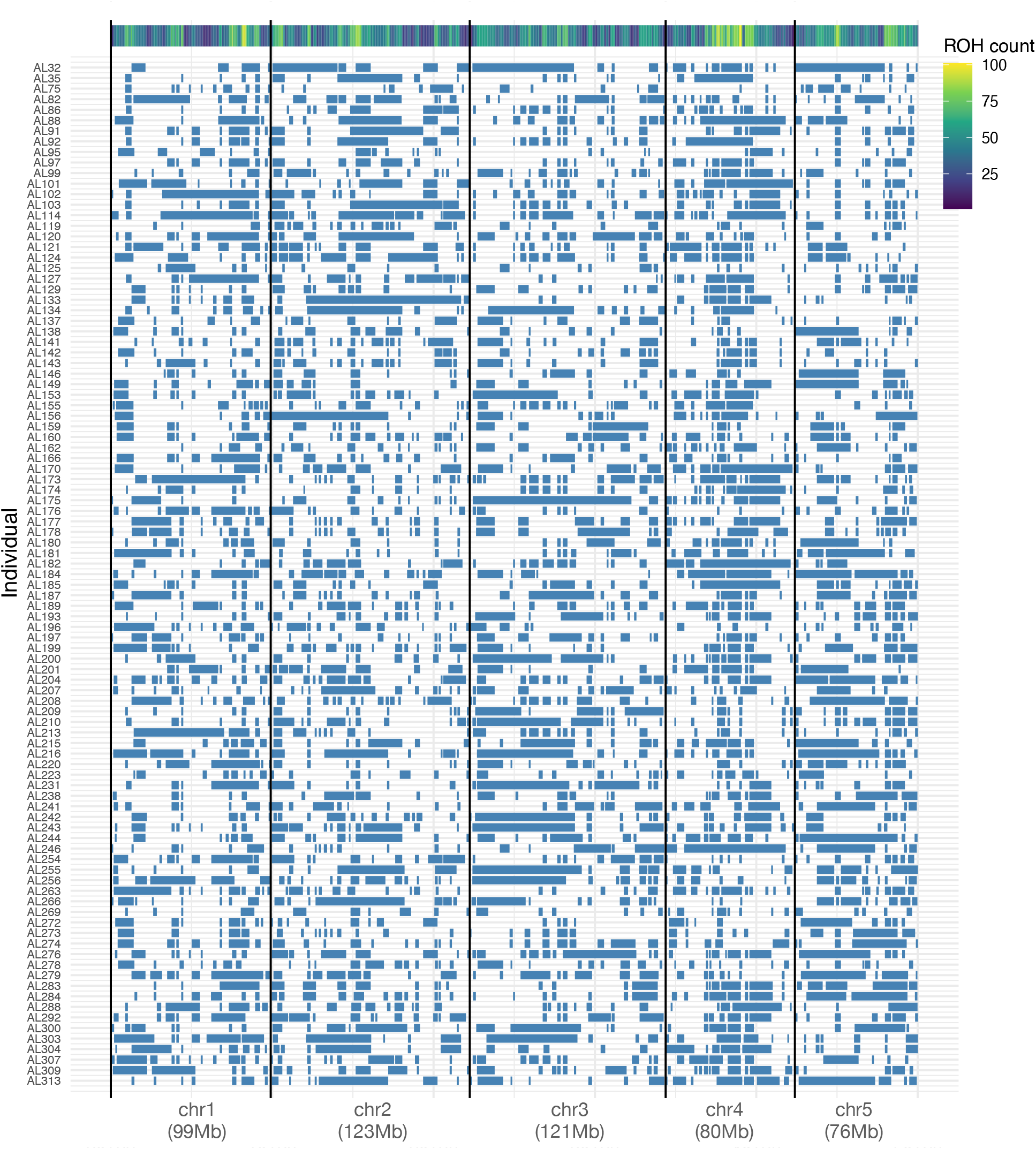
Runs of homozygosity visualized across the first five chromosomes for all 97 hatched ’Alalā individuals. Heatmap of ROH abundance shown on the top. See Figure S1 for the same plot for all 78 embryos that failed to hatch.

**Figure 2:**
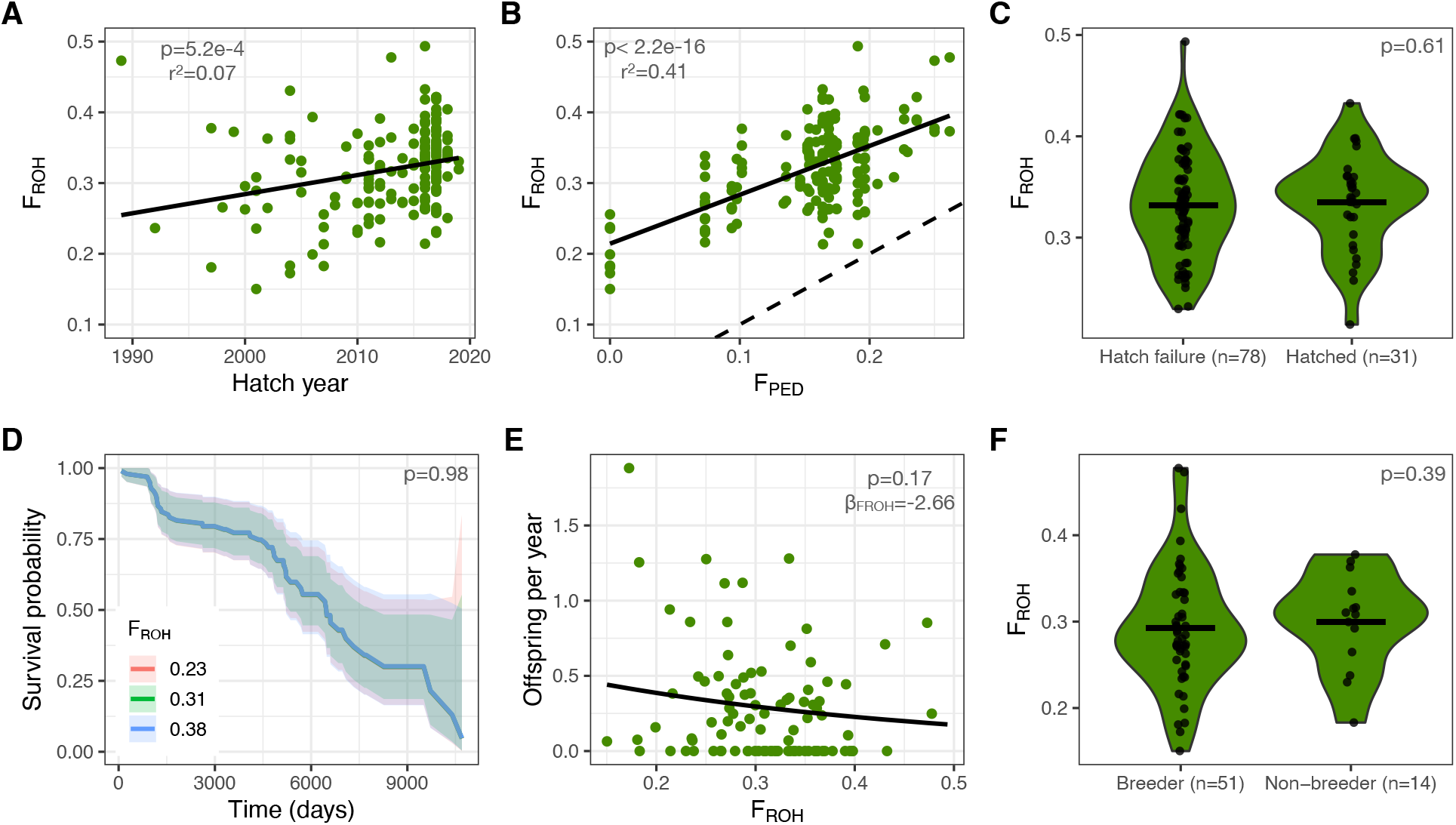
Patterns of inbreeding and inbreeding depression in the ’Alalā conservation breeding program. (A) Levels of inbreeding (measured by F_ROH_ for ROH > 1Mb) increase over time in hatched individuals (n=97), though are elevated even in the 1990s. (B) F_ROH_ is correlated with, though much greater than, F_ped_. Dashed line represents 1:1 relationship. (C) Levels of F_ROH_ do not differ between 78 embryos that failed to hatch and 31 hatched individuals born during the same time period. (D) No impact of F_ROH_ on adult survival in a Cox proportional hazards model (p=0.98). See Table S2 for full model output. (E) F_ROH_ is negatively correlated with annual reproductive output, though this relationship is not statistically significant. See Table S3 for full model output. (F) No differences in F_ROH_ between 51 individuals that produced viable offspring at any point in their life versus 14 individuals that never successfully reproduced. Note that horizontal black lines in panels C and F denote the means of each distribution. See Figures S3 and S4 for the same results with an ROH threshold of 100kb and 10Mb and Figure S6 for results when excluding low coverage individuals.

In part due to this high abundance of ROH, overall levels of genetic diversity in ‘Alalā are very low (average observed heterozygosity = 0.46 hets/kb), among the lowest observed in birds (**Fig. S5**). Low genetic diversity in ‘Alalā is also a consequence of relatively low levels of heterozygosity in non-ROH regions (0.67 hets/kb) due to modest historical effective population sizes on the order of ∼35,000 (**Fig. S7**). When examining spatial patterns of nucleotide diversity across 50 individuals alive in 2025, we observe very few regions of the genome where genetic diversity has been entirely lost (**Fig. S8**). Thus, although genetic diversity is substantially reduced in the present-day ‘Alalā population, few ROH are fixed and some degree of haplotype diversity has been preserved.

### Limited impacts of genomic inbreeding on survival and reproduction

We next tested for associations between F_ROH_ and measures of survival and reproductive success. In particular, we were interested in contrasting F_ROH_ between embryos that failed to hatch and individuals that successfully hatched, given previous evidence suggesting that inbreeding may be responsible for the low hatching success rate in ‘Alalā ^20,21^. In contrast to these studies, we found no evidence to suggest that genome-wide inbreeding was associated with embryo mortality. Specifically, we observed no differences in F_ROH_ when comparing 78 embryos that died in egg and 31 individuals that successfully hatched during the same time period (Wilcoxon rank sum test p=0.61; **Fig. 2C**). Similarly, when running Cox proportional hazards models, we found that F_ROH_ was not statistically associated with adult survival (n=97; p=0.98; **Table S2**), exhibiting identical survival curves for low (0.23) and high (0.38) F_ROH_ quantiles (**Fig. 2D**). Overall, these results suggest that the impact of autozygosity on hatching success and adult survival in ‘Alalā is weak and not detectable even with relatively large sample sizes.

When testing for correlations between F_ROH_ and reproduction, we similarly found little evidence for ROH driving reduced fitness. We ran negative binomial models relating nestling production and parent F_ROH_ while accounting for hatch year and longevity, finding a negative though non-significant (p=0.17) relationship between F_ROH_ and the number of nestlings produced per year (**Fig. 2E; Table S3**). Because mate pairing opportunities in the ‘Alalā breeding program have varied over time and could confound these results, we examined a potentially more robust contrast of F_ROH_ between individuals >10 years of age that successfully reproduced at any point in their lifetime (n=42) and those that did not (n=10). Here again, we found no evidence for an association between F_ROH_ and reproduction (**Fig. 2F**). Finally, we also tested for correlations between F_ROH_ and the average annual egg laying rate and egg hatching rate across the lifetime of an individual, observing no statistically significant associations (**Fig. S9**).

To explore the sensitivity of our results to the ROH length threshold, we reran the above analyses with both shorter (100 kb) and longer (10 Mb) thresholds. As above, we found no statistically significant associations between any survival or reproductive metrics and F_ROH_ (**Figs. S3 & S4**). Finally, we also explored the impact of variable sequencing depth by removing low coverage (<10x) individuals, again finding no significant correlations between F_ROH_ and any fitness component (**Fig. S6**).

### Prevalent recessive lethal alleles contribute to hatching failure

Although the overall ROH burden does not appear to be associated with hatching failure in ‘Alalā, it is possible that specific genomic variants may still be contributing to low hatching success. To test for variants associated with hatching failure, we conducted a genome-wide association analysis using a recessive genetic model, identifying SNPs where the homozygous alternate genotype was disproportionately enriched among embryos that failed to hatch relative to hatched individuals. Two regions on chromosomes 10 and 13 stood out with notable associations (**Fig. 3A**).

**Figure 3:**
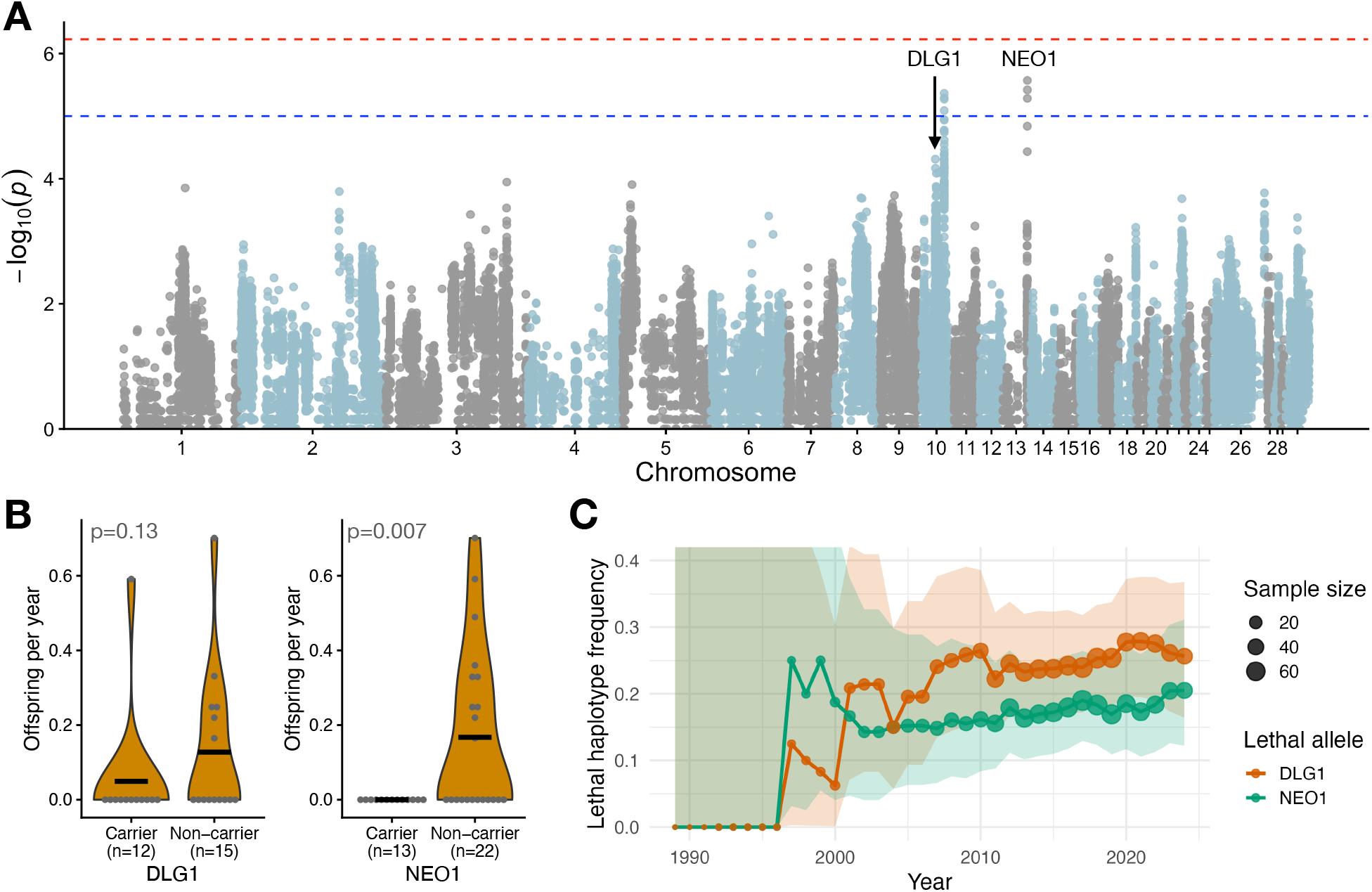
Identification of two recessive lethal variants in essential developmental genes on chromosomes 10 (DLG1) and 13 (NEO1). (A) Results from an association analysis specifically testing for recessive lethal alleles associated with egg hatching failure. The dotted red line represents the Bonferroni-corrected p-value threshold of p=5.9e-7 and the dashed blue line represents a suggestive threshold of p=1e-5. See Figure S10 for a zoomed in plot of the two adjacent peaks on chromosome 10. (B) Reproductive rates are lowered in heterozygous carriers at each recessive lethal locus compared to non-carriers for sequenced individuals born in the breeding program after 2012. Means shown with horizontal black bar. See Figure S12 for full results. (C) Recessive lethal allele frequencies plotted from all hatched individuals (n=97). Allele frequencies for each year computed from individuals living during that year, with the dot size corresponding to the sample size.

On chromosome 10, we detected six suggestive SNPs (p<1e-5) in a region annotated as a pseudogene, in high linkage disequilibrium (LD; r^2^=0.75) with another region ∼6.5 Mb upstream that also contains several associated SNPs (p<1e-4) within protein coding genes (**Figs. 3A & S10**). These substitutions included a missense SNP in DLG1 from Cys → Tyr at position 23/766 predicted to have moderate deleteriousness, where 8/8 individuals that were homozygous for the alternate allele failed to hatch. Hatching rates for heterozygous (n=65) and homozygous reference (n=77) genotypes were both near 60% (**Fig. S11**). DLG1 is a compelling candidate for contributing to embryonic lethality, as it is a key developmental gene that is expressed in chicken embryos and is lethal when knocked out in mice ^29–31^. Across this region of chromosome 10, the observed patterns of LD and coverage are consistent with a ∼3.5 Mb inversion polymorphism (**Fig. S10**).

On chromosome 13, we detected three SNPs spanning 36 kb that are strongly associated with hatching failure (p<1e-5), with 15/15 homozygous alternate individuals failing to hatch (**Fig. 3A**). Here again, hatching success rates for heterozygous (n=53) and homozygous reference (n=99) individuals were both near 60% (**Fig. S11**). These variants are splice-region SNPs affecting exon–intron boundaries in NEO1, a key developmental gene known to cause cell apoptosis when overexpressed in chicken embryos and to be embryonic lethal when knocked out in mice ^32–34^. Taken together, these findings suggest that these variants could potentially be impacting the expression of NEO1 and causing cell apoptosis in developing ‘Alalā embryos, though more work is needed to confirm this functional link.

To explore the fitness effects of these variants in heterozygous carriers, we contrasted annual offspring production between carriers vs non-carriers for individuals born after 2012 (see **Methods**). Strikingly, we find that only one out of 25 heterozygous carriers across both loci ever successfully reproduced (4%), compared to 16/37 (43%) non-carriers (**Figs. 3B & S12**). These differences in reproductive rates between carriers and non-carriers were statistically significant at the NEO1 locus (Wilcoxon p=0.007), though not at the DLG1 locus (p=0.13). Overall, these findings further highlight the substantial impacts of these recessive lethal loci on reproductive fitness in ‘Alalā.

Despite their large effect on reproductive fitness, these recessive lethal alleles remained surprisingly abundant in the ‘Alalā breeding program. Specifically, lethal allele frequencies derived from all 97 sequenced hatched individuals were typically around 25% over the last 20 years for DLG1 and between 15-20% for NEO1 (**Figs. 3C & S13**). Notably, we do not observe evidence of purifying selection at either locus decreasing the recessive lethal allele frequencies over time, suggesting that, despite the large effects of these alleles, they continue to persist in the population. However, our ability to more fully investigate allele frequency trajectories remains limited by a very small number of sequenced individuals born prior to 2000 (n=6), as well as no sequenced individuals born after 2018 (**Table S1**).

Finally, we also note that, although homozygous recessive lethal genotypes are found exclusively within long ROH at the DLG1 locus on chromosome 10, the same does not appear to be true at the NEO1 locus on chromosome 13 (**Fig. S14**). Thus, these recessive lethal variants could potentially be overlooked when exclusively focusing on long ROH.

### Eco-evolutionary simulations recapitulate impacts of the population bottleneck

To understand whether our empirical findings of weak effects of ROH combined with an outsized impact of recessive lethals are plausible under a set of reasonable demographic and genomic parameters, we parameterized an eco-evolutionary Wright-Fisher model for ‘Alalā in SLiM ^35,36^. We assumed a mutation rate typical for passerine birds ^37–39^, selection and dominance parameters that have previously been validated for vertebrates ^16,40^, a historical effective population size of 33,500, and life history parameters informed by the captive population (see **Methods**). For each simulation replicate, we modeled the initial establishment of the captive breeding population from 1973-1993, during which the wild population was declining towards extinction ^41^ and allowed the breeding program to grow to its present-day size of ∼120 individuals.

With this model, we first explored how the population bottleneck and subsequent recovery is expected to impact F_ROH_, fitness, and inbreeding load (i.e., the diploid number of lethal equivalents, or 2B ^40,42^), and how these quantities may be expected to change moving into the future (**Fig. 4A**). We observe that F_ROH_ values rapidly increase from 1973 to present day in close agreement with empirical values, and are predicted to continue to increase to F_ROH_=0.39 by 2100 (**Fig. 4A**). Due to this accumulation of inbreeding, fitness declines from an initial value of 0.967 to a minimum of 0.949 in 2007 before recovering to 0.965 in 2100 (**Fig. 4A**). Notably, this near-complete recovery of fitness suggests a minimal impact of fixed deleterious mutations, likely due to the relatively rapid recovery from the initial bottleneck in just ∼3 generations. Finally, the predicted inbreeding load for survival to age 6 is substantially purged from 2B=2.8 in 1975 to 2B=1.9 in present day to 2B=1.3 in 2100, consistent with ongoing purging of recessive deleterious variants that are subject to greater purifying selection after being exposed by inbreeding. Notably, the initial magnitude of the inbreeding load is lower than a median empirical estimate for survival to sexual maturity from wild vertebrates of 2B=4.5 ^43^, suggesting that inbreeding depression in ‘Alalā was already somewhat weak prior to the bottleneck, and has been further diminished by purging.

**Figure 4:**
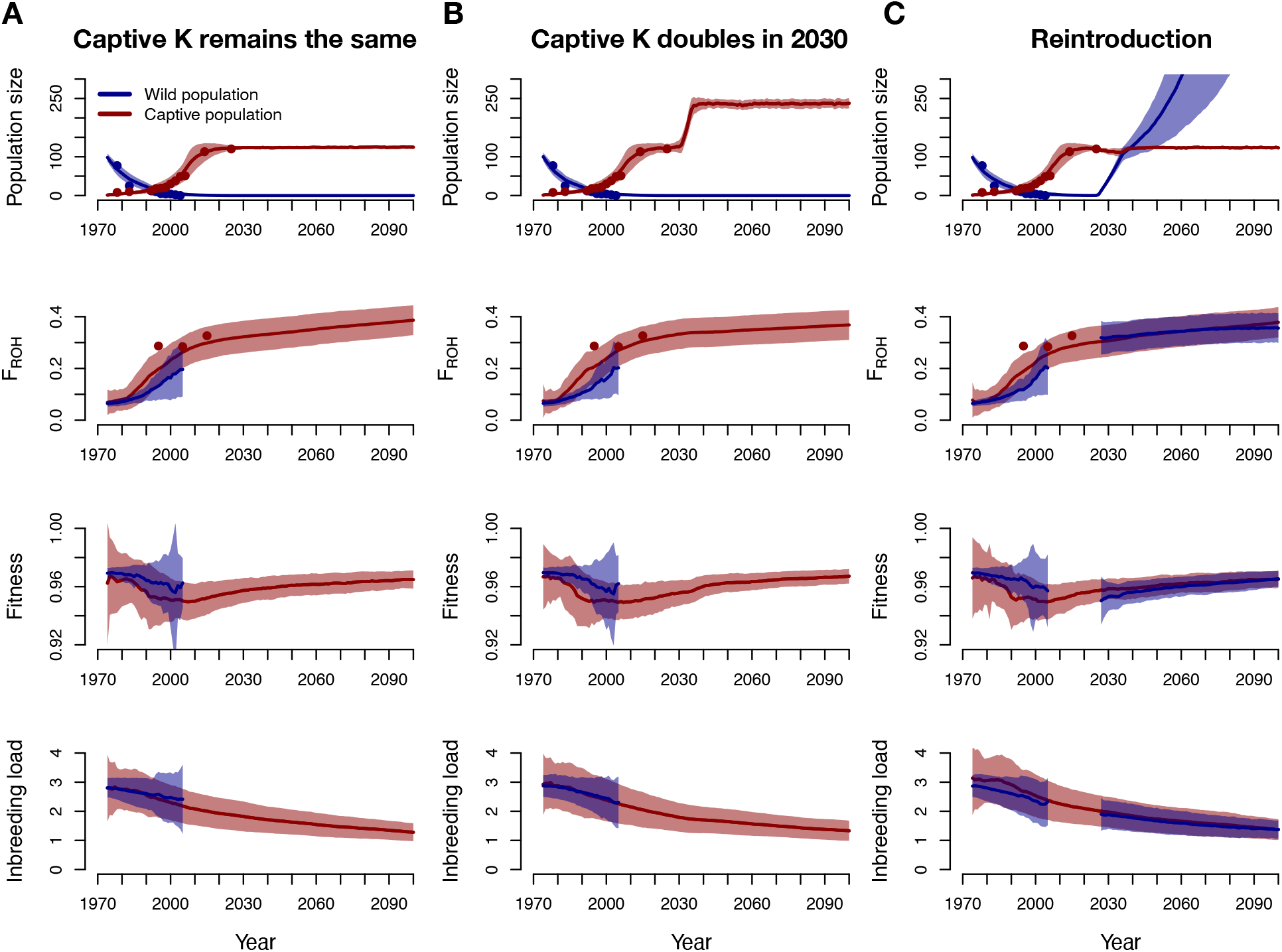
Eco-evolutionary simulation results under three hypothetical management strategies. Captive population depicted in red, wild population in blue, and points represent empirical census estimates. (A) Carrying capacity for the captive population remains the same (K=130) moving into the future. (B) Carrying capacity for the captive population increased to K=260 by 2030. (C) Reintroduction of 10 individuals per year from 2026-2036 assuming an elevated mortality rate in the wild of 10%. For each simulated scenario, the top row shows projected population sizes, the second row shows projected F_ROH_, the third row shows projected fitness, and the bottom row shows projected diploid inbreeding load (2B). Results shown from the founding of the captive population in 1973 projected into the future by 2100, with shading showing one standard deviation.

To directly test whether this model could recapitulate our empirical findings, we examined the relationship between F_ROH_ and reproduction from the simulation output. We observe a negative but non-significant relationship between F_ROH_ and nestlings per year (**Fig. 5A**) that is strikingly similar to our empirical results (**Fig. 2E**), suggesting that weak inbreeding depression is theoretically expected for ‘Alalā as a consequence of low non-ROH heterozygosity. In other words, because ‘Alalā naturally harbors relatively low levels of segregating genetic variation, there are few heterozygous recessive deleterious alleles that can be unmasked by inbreeding and contribute to inbreeding depression. Indeed, when we increase the expected non-ROH heterozygosity in this model three-fold from 0.67 to 2.0 hets/kb (see **Methods**), we observe a much greater simulated pre-bottleneck inbreeding load (2B≈10 vs 2B≈3) which translates to a stronger and statistically significant relationship between F_ROH_ and reproduction (**Fig. 5B**). Notably, these higher levels of non-ROH heterozygosity of 2.0 hets/kb closely mirror that of ‘akikiki, another critically endangered Hawaiian bird subject to similar management for which strong correlations between ROH and reproduction have previously been reported ^16^.

**Figure 5:**
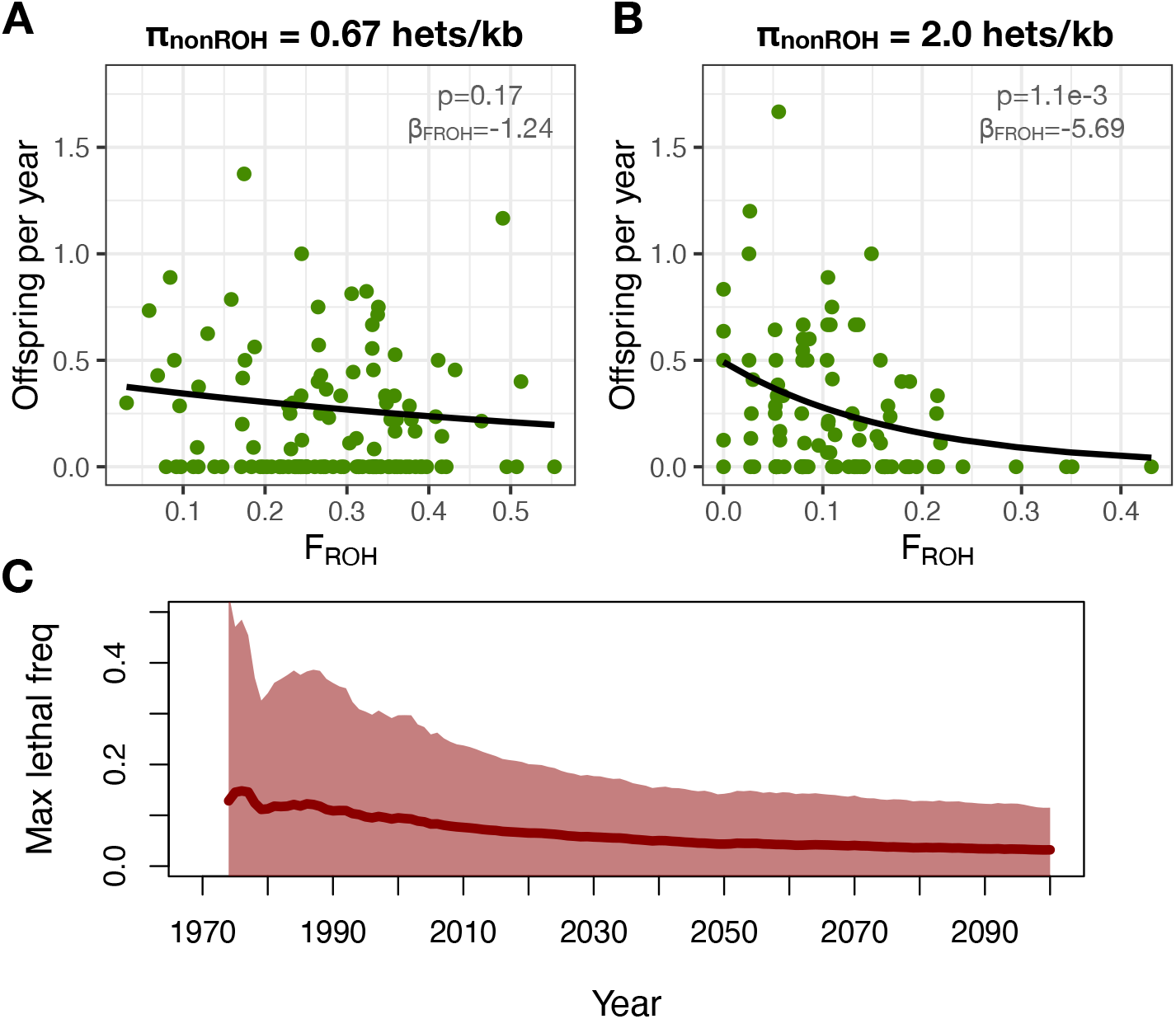
Theoretical predictions mirror empirical results in ‘Alalā. (A) A negative but not statistically significant relationship between F_ROH_ and annual reproductive output is predicted from a model with the empirical level of non-ROH heterozygosity observed in ’Alalā (0.67 hets/kb). (B) A much stronger and statistically significant relationship between F_ROH_ and annual reproductive output is predicted from a model with 3x increased levels of non-ROH heterozygosity (2.0 hets/kb). (C) Predicted maximum recessive lethal allele frequencies during and after the population bottleneck. Average maximum recessive lethal frequency shown with dark red line and shading shows two standard deviations. Note that high recessive lethal allele frequencies >0.2 are observed during and after the founding bottleneck, though are attenuated after ∼2020.

To explore whether a high abundance of recessive lethals (**Fig. 3C**) is also predicted by our model, we plotted the maximum recessive lethal allele frequency during the simulated bottleneck and recovery period (**Fig. 5C**). During the bottleneck, maximum lethal allele frequencies increase to an average of ∼15% and often higher than 30% (**Fig. 5C**), compared to <1% frequency prior to decline (**Fig. S15**). However, these extreme allele frequencies become attenuated after the population recovers to N>100 and purifying selection becomes more effective, eventually reducing to ∼5% by 2100 (**Fig. 5C**). Thus, even as many highly deleterious alleles are lost during the bottleneck, some may drift to high frequency and potentially result in severe fitness consequences.

### A path to recovery for ‘Alalā

We next explored several proposed management scenarios for ‘Alalā, quantifying predicted effects on genomic erosion. First, we investigated the impacts of increasing the carrying capacity of the breeding program, given concerns that continuing to maintain the captive population at ∼120 could further bottleneck the species. When modelling a hypothetical doubling of the captive carrying capacity in 2030, we observe only modest differences in predicted F_ROH_ (0.368 vs 0.386) and fitness (0.967 vs 0.965) in 2100 compared to a scenario where the carrying capacity remains unchanged (**Fig. 4**). These modest differences are due in part to the relatively small number of generations (∼11) that transpire between 2030 and 2100. Overall, these findings suggest that doubling the number of available aviaries may only have modest impacts on inbreeding and deleterious genetic load over the relatively short term of ∼70 years.

The ultimate goal for ‘Alalā conservation is to recover a wild population. However, initial and limited efforts to release individuals have been plagued by high mortality in the wild, likely due to limited food resources ^44^. To test whether captive breeding and release can overcome this mortality to establish a viable wild population, we modeled release scenarios varying cohort sizes and mortality rates. When releasing 100 individuals over ten years (2026-2036), lower wild mortality rates led to faster population recovery, though recovery failed in the highest mortality scenario (15% above captive rates; **Figs. 5 & S16**). Releasing fewer individuals (50 total) also prevented recovery, while releasing 150 resulted in a more rapid and robust recovery (**Fig. S17**). Encouragingly, several scenarios produced rapid wild population recovery, particularly when mortality rates are low and ≥100 individuals are released. In these scenarios, F_ROH_ declines in the wild relative to the captive population and fitness increases, suggesting a path to demographic and genomic recovery.

## Discussion

In this study, we present a comprehensive analysis of inbreeding depression in ‘Alalā, a species that recovered from the brink of extinction but has yet to recover in the wild. Despite a severe population bottleneck and a high abundance of long ROH, we find little evidence that genome-wide inbreeding predicts variation in survival or reproduction, including hatching success. Instead, we identify two recessive lethal haplotypes segregating at high frequency that together explain nearly all of the presumed genetic contributions to hatching failure. Our findings reveal a decoupling between ROH and fitness in this system and suggest that the fitness consequences of severe founder events may be driven less by genome-wide inbreeding and more by the stochastic rise of a small number of recessive lethal alleles.

Multiple lines of evidence suggest the variants identified at DLG1 and NEO1 have large and measurable impacts on fitness, exhibited by 100% hatching failure for homozygous individuals and severely reduced reproductive success in heterozygous carriers. Notably, the present-day allele frequencies of these recessive lethal variants (∼25% and ∼20%) imply homozygous lethal genotype frequencies of ∼6.25% and ∼4%, respectively, or a total hatching failure burden of ∼10%. Thus, these two recessive lethal regions alone explain nearly all of the excess hatching failure in ‘Alalā (>50%^21^) compared to the baseline rate for captive bird species (∼38%^23^). Although more work is needed to detail the functional pathways impacted by putative lethal variants in DLG1 and NEO1, both of these critical developmental genes are expressed in chicken embryos and associated with embryonic lethality in mice, further supporting their potential impacts on hatching failure in ‘Alalā ^29,30,32–34^.

A key takeaway from our work is that, although species can recover from severe bottlenecks despite high inbreeding, this may come at the cost of elevated frequencies of recessive lethal alleles. Although most recessive lethal mutations are expected to be purged during population bottlenecks, the elevated frequency of one or a few of these alleles via founder effects can nevertheless result in severe fitness consequences. Similar patterns observed in other bottlenecked species ^49–51^ suggest that elevated recessive lethal frequencies may be a broadly important consequence of population decline. Notably, although previous studies have emphasized the accumulation of weakly deleterious mutations as the primary genetic consequence of bottlenecks ^45–48^, our findings suggest a limited impact of these variants on fitness in ‘Alalā compared to the outsized impact of recessive lethals.

An optimistic outlook for ‘Alalā is that these recessive lethal alleles can still be purged in the future. However, although our simulation results predict a decline in lethal allele frequencies, empirical allele frequency trajectories do not exhibit this expected decline, perhaps suggesting the presence of linkage leading to selective interference ^52^, balancing selection ^53^, or reproductive compensation ^54^. In the captive environment specifically, reproductive compensation may be particularly pronounced: managers routinely provide additional breeding opportunities to pairs with poor hatching success, potentially offsetting the fitness cost of carrying lethal alleles and slowing their removal from the population. Additionally, the presence of a putative inversion polymorphism near DLG1 could suppress recombination and magnify the effects of selective interference; such impacts of inversions resulting in high frequency lethals have been extensively documented in *Drosophila* ^52^. Future work with expanded temporal and long read genomic datasets is needed to better map this putative inversion and more fully investigate the function and frequency of these recessive lethal loci in ‘Alalā.

Our findings contrast with mounting evidence from other species that ROH are often strongly associated with reduced fitness ^14,55,56^, as well as previous studies reporting impacts of pedigree-based inbreeding on survival in ‘Alalā ^20,21^. Differences from previous studies of ’Alalā may reflect the fact that F_ped_ and F_ROH_ measure related but distinct aspects of inbreeding history and that previous studies analyzed survival across broader life history stages using different modelling approaches ^20,21^. Notably, our simulation analysis demonstrates that these limited associations between ROH and fitness are theoretically expected for ‘Alalā due to low non-ROH heterozygosity contributing to a low inbreeding load ^11,14,57^. The agreement between our empirical and simulation-based relationship between F_ROH_ and reproduction is particularly striking given that our simulation model was parameterized only with fundamental demographic, genomic, and life history parameters for the species, and without any direct fitness data. This finding therefore suggests that, in cases where direct measures of inbreeding depression are unavailable, genomics-informed simulation models can provide a useful tool for exploring the relationship between deleterious genetic variation and fitness ^40,47,58,59^.

Together, our results carry several implications for the management of ‘Alalā and similarly bottlenecked species. Our identification of two recessive lethal loci modulating hatching failure in ‘Alalā highlights the possibility of increasing egg hatching rates by reducing the frequency of these lethal haplotypes. In practice, this could be achieved through genotype-informed pairing strategies that avoid carrier × carrier matings and by prioritizing individuals with lower lethal carrier burden for wild release, limiting the introduction of these alleles into the wild population. As reintroduction efforts continue, such genomic benchmarks may ultimately inform longer-term recovery milestones and exit criteria for intensive management – helping to define the conditions under which ’Alalā can thrive in the wild.

## Acknowledgements

Funding was provided by the Royal Society of New Zealand’s Marsden Fund project ‘Resolving the genomic architecture of hatching failure to improve conservation of endangered birds’ (Contract ID UOO1817) awarded to BCR, JTS, OAR and CEG, and Howard Hughes Medical Institute (HHMI) awarded to EDJ. New Zealand’s national facilities are provided by NeSI and are funded jointly by NeSI’s collaborating institutions and the Ministry of Business, Innovation and Employment (MBIE) research infrastructure programme. EBC was supported through the UH Hilo Pacific Islands Program for Exploring Sciences (PIPES) via funding from the National Science Foundation (NSF, Award # 2447178). The Biodiversity Banking team of San Diego Zoo Wildlife Alliance established the ’Alalā fibroblast cell line and karyotype.

## Author Contributions

Project conception: CCK, SG, YF, BM, OAR, CEG, JTS, BR, APW

Reference genome assembly: GF, JB, AT, JMDW, SG, BM, ED, OF, EDJ, JTS, OAR, BR

Short read data generation: SG, ED, BM, JTS, OAR, BR

Computational analysis: CCK, SG, YF, BM, AMF, JAR, EBC, APW

Manuscript drafting: CCK

Manuscript editing: All authors

## Data and Code Availability

The ’Alalā reference genome (bCorHaw1.pri.cur) is available on NCBI under GenBank ID GCA_020740725.1 and raw sequence data is available from the Sequence Read Archive under BioProject PRJNA1499171. All analysis scripts are available on GitHub: https://github.com/ckyriazis/alala_genomics.

## Methods

### Reference genome assembly and annotation

A chromosome-scale reference genome was generated for a female individual AL308 from the captive breeding program (born July 5, 2018), using the Vertebrate Genomes Project (VGP) pipeline. A female was chosen to obtain both Z and W sex chromosomes. Whole blood samples were used for library generation and sequencing, including four 100 µL aliquots preserved in ethanol. They were separated in 100 µL aliquots preserved in RNA/DNA Shield for transcriptome sequencing (although genome sequencing with long reads works better without such preservatives). High molecular weight DNA > 300 Kb was extracted and used for PacBio HiFi sequencing (∼30x coverage, ∼20 Kb inserts), Bionano optical mapping (∼100x), and Hi-C sequencing (∼60x), followed by genome assembly and curation. Genome assembly was performed using the VGP pipeline v2.0 implemented on Galaxy ^24^. Transcriptome data for genome annotation were generated using VGP Iso-Seq protocols from whole blood, including RNA extraction, quality control, Iso-Seq library preparation, and sequencing on one PacBio Sequel II SMRT Cell. Manual curation was performed to fix structural errors made in the automated assembly process, remove non-crow contaminating genomic sequence (common for all assemblies), and to name chromosomes. The mitochondrial genome was assembled using mitoVGP ^60^.

Whole genome alignments were generated between the new chromosome-scale ’Alalā genome and three other corvid chromosome-level genomes: hooded crow (*C. cornix cornix*, GCA_000738735.5 ^61^), New Caledonian crow (*C. monedoloides*, GCA_009650955.1 ^62^), and Eurasian jackdaw (*Coloeus monedula*, GCA_013407035.1 ^63^). Pairwise alignments were computed using LastZ v1.04 ^64^ and synteny visualised with Circos v0.69 ^65^. These alignments helped to name chromosomes as consistently as possible.

The genome was annotated using the NCBI Eukaryotic Genome Annotation Pipeline (EGAP), v100, by NCBI. Long read Pacbio IsoSeq data as well as other avian transcriptome data in NCBI was used for the annotation. Transcriptome data available in NCBI from four other crow species was used for the annotation (*C. corone*, *C. cornix*, *C. woodfordi*, and *C. kubaryi*), totalling over 2.8 billion reads.

### DNA extraction and population resequencing

Genomic DNA was extracted from samples of 175 individuals (whole blood from adults and frozen embryos) using a standard phenol-chloroform extraction protocol. For blood samples, 5-20 µl of blood was used depending on the dilution of the mixture. Frozen embryos were carefully thawed on ice and a small piece of tissue or the entire embryo (for early death) was cut with a sterilized scalpel and then ground into a lysis buffer with a pestle until dissolved.

Genomic DNA was sequenced at the UC Davis DNA Technologies & Expression Analysis Core Laboratory. Barcode-indexed sequencing libraries with an insert size of 330-380 bp were generated from genomic DNA samples sheared on an E220 Focused Ultrasonicator (Covaris, Woburn, MA). For each sample 200 ng of sheared DNA was converted to sequencing libraries using a Kapa High Throughput Library Preparation Kit (Kapa Biosystems-Roche, Basel, Switzerland). The libraries were amplified with 4 and 8 PCR cycles and analyzed with a Bioanalyzer 2100 instrument (Agilent, Santa Clara, CA), quantified by fluorometry on a Qubit instrument (LifeTechnologies, Carlsbad, CA), and combined into one pool at equimolar ratios. The pool was quantified by qPCR with a Kapa Library Quant kit (Kapa Biosystems-Roche) before sequencing on a NovaSeq 6000 sequencer (Illumina, San Diego, CA) with 150 bp paired-end reads.

### Short-read population data processing

We processed short-read sequence data using a pipeline adapted from the Genome Analysis Toolkit (GATK) Best Practices Guide ^66^. First, we trimmed adapter sequences using CutAdapt ^67^ and aligned trimmed sequences to the reference genome using BWA-MEM ^68^. We then used sambamba ^69^ to mark duplicate reads and assess coverage and alignment statistics for each sample. We used HaplotypeCaller in GATK v3.8 to perform genotype calling at all sites in each sample individually and then performed joint genotyping across all samples using GenotypeGVCFs.

We filtered genotypes to include only high-quality biallelic SNPs and monomorphic sites, removing sites with Phred score below 30 and depth exceeding the 99th percentile of total depth across samples. In addition, we removed sites that failed slightly modified GATK hard filtering recommendations (QD < 4.0 || FS > 12.0 || MQ < 40.0 || MQRankSum< − 12.5 || ReadPosRankSum < − 8.0 || SOR >3), as well as those with >25% of genotypes missing or >75% of genotypes heterozygous and those falling within repetitive regions as identified using Tandem Repeats Finder ^70^ and WindowMasker ^71^. At an individual level, we also removed sites with less than 3x coverage or with coverage exceeding 2.5 times the individual mean. We restricted all downstream analysis to the 41 assembled autosomes, together comprising 1.04 Gb in length. After filtering, samples had 18.3x read depth on average (range: 1.03-87.2; **Table S1**).

### Genomic diversity, runs of homozygosity, and historical demography

We estimated observed heterozygosity for all 175 sequenced individuals by dividing the number of heterozygous sites by the total number of called sites (variant and invariant) using a custom R script ^72^. To assess the extent to which genetic diversity may have been lost in the current ’Alalā population due to fixed haplotypes, we plotted patterns of nucleotide diversity across the genome from 50 individuals alive in 2025 in 100 kb windows inferred using pixy ^73^. We compared levels of genetic diversity in the current ’Alalā population to other avian taxa by plotting the average observed heterozygosity in ’Alalā along with estimates from 41 bird species following Kyriazis et al. ^16^. As heterozygosity is strongly correlated with sequencing depth (**Fig. S18**), we estimated observed heterozygosity only from high-coverage (>20x) living individuals (n=16). Note that average observed heterozygosity estimates from high coverage individuals and genome-wide nucleotide diversity estimates from pixy were highly similar (4.73e-4 vs 4.75e-4).

We inferred runs of homozygosity using BCFtools/ROH ^74^ using the -G30 flag and visualized ROH along the first five chromosomes of all sequenced individuals using a custom R script. To estimate the inbreeding coefficient (F_ROH_) for each individual, we calculated the proportion of the autosomal genome covered by ROH >1Mb and >10Mb in length. We plotted the relationship between F_ROH_ and hatch year and between F_ROH_ and pedigree-based F (F_ped_) estimated using PMx ^75^ . Note that, unlike heterozygosity, F_ROH_ was not correlated with sequencing depth (**Fig. S18**).

To estimate non-ROH heterozygosity, we divided the average heterozygosity from 61 high coverage (>20x) individuals (0.455 hets/kb) by 1-F_ROH_. When assuming an ROH threshold of 1Mb, average F_ROH_=0.319, yielding non-ROH heterozygosity of 0.67 hets/kb. However, when assuming a shorter ROH threshold of 100kb, average F_ROH_=0.395, resulting in a slightly higher non-ROH heterozygosity of 0.75 hets/kb.

To infer the long-term demographic history of ’Alalā, we employed the Pairwise Sequential Markovian Coalescent (PSMC) model ^76^, which estimates historical effective population size trajectories from patterns of heterozygosity in the diploid genome of a single individual. Filtered VCF files were converted to consensus FASTA files, which were then used to generate PSMC input files using the fq2psmcfa function. For presenting our findings, we selected the oldest individual from our dataset (AL032, born in 1989) to minimize impacts of the recent bottleneck, though repeated this analysis with 10 total individuals to confirm consistency of results. Based on recommendations from Hilgers et al. ^77^ we set the time intervals using the -p flag as 2+2+25*2+4+6. As elsewhere, we assumed a mutation rate of 5e-9 mutations per site per generation ^37–39^. Based on our simulation analysis, we assumed a generation time of 6 years (see below). To contextualize inferred effective population sizes, we plotted results alongside Ne estimates for three Hawaiian honeycreeper species from Kyriazis et al. ^16^.

### Testing for correlations between F_ROH_ and fitness

To investigate for potential associations between ROH and egg hatching failure, we contrasted F_ROH_ between individuals that hatched and those that failed to hatch. An initial comparison of all hatched (n=97) versus non-hatched (n=78) individuals suggested that non-hatched individuals had higher F_ROH_ (mean F_ROH_ = 0.33 for non-hatched vs 0.31 for hatched, Wilcoxon p=0.016). However, hatch years systematically differ in our sample between birds that hatched (1989-2018) and embryos that failed to hatch (2014-2019). Because embryos that failed to hatch were only sampled in the last five years of the 30-year study, hatching success may be confounded with the overall increase in F_ROH_ over time in the breeding program (**Fig. 2A**). To account for this, we subsetted our sample of hatched individuals to include only individuals born after 2014 (n=31).

We tested for correlations between F_ROH_ and adult survival in our complete sample of hatched individuals (n=97) using a Cox proportional hazards model with F_ROH_ included as a fixed effect. Observations of survival of birds still living were right-censored to account for the incomplete observation. Models were fitted using the survival package in R ^72^. Model fit was visualized by plotting survival curves for F_ROH_ quantiles of 10%, 50%, and 90%.

To test for associations between F_ROH_ and lifetime reproductive success in our complete sample of hatched individuals (n=97), we used a negative binomial generalized linear mixed model with a log link. Total offspring number was modeled with an offset for lifespan. Hatch year of the breeding individual was included as a random intercept to account for cohort effects, and F_ROH_ was included as a fixed effect. Models were fit using the glmmTMB in R ^72^. Because opportunities for reproduction vary over time in the breeding program, we also explored a potentially more robust comparison of F_ROH_ between individuals that successfully bred at any point and those that did not (restricting to individuals >10 years of age that had ample opportunities to reproduce, n=65). Finally, we also tested for correlations between F_ROH_ and annual rate of laying eggs and egg hatching rate averaged across breeding seasons in all hatched individuals (n=97) using simple linear regression.

### Identifying recessive lethal variants underlying hatching failure

We tested for variants associated with egg hatching failure using an association analysis specifically aimed at identifying embryonic recessive lethal mutations. For each of 1,111,422 autosomal SNPs, we tested a recessive genetic model in which individuals homozygous for the alternate allele were contrasted against all other genotypes (heterozygotes and homozygous reference). This framework can detect variants across a spectrum of recessivity, as long as homozygous alternate individuals have a higher probability of hatching failure than the combined heterozygous and homozygous reference classes. Association testing was performed using Fisher’s exact tests on 2x2 contingency tables, comparing the frequency of homozygous alternate genotypes between embryos (n=78) that died prior to hatching and those that hatched successfully (n=97). To focus specifically on putative recessive lethal variants, SNPs were tested only if the homozygous alternate genotype was enriched among embryos that died and rare among hatched individuals (frequency <5% in hatched individuals and >5% in embryos that died). We plotted -log10(p-values) alongside a suggestive p-value threshold of p<1e-5 and a Bonferroni significance threshold of p<5.9e-7.

Because the objective of this analysis was to identify candidate recessive lethal variants segregating within family groups rather than to estimate marginal SNP effects across an unrelated population, we did not adjust for relatedness or population structure. Recessive lethal alleles are expected to reach homozygosity primarily through recent shared ancestry (i.e., inbreeding), and correcting for kinship would remove biologically meaningful signal rather than confounding. Statistical tests were therefore used as a heuristic to rank candidate loci showing enrichment of homozygous alternate genotypes among embryos that failed to hatch, rather than as formal tests of genome-wide significance.

To assess the potential functional consequences of candidate variants underlying the two strongest association signals, we annotated all sequence variants within the top associated regions on chromosomes 10 and 13 using snpEff v5 ^78^. We built a custom annotation database using our annotated gene models and classified variants by gene context (intergenic, intronic, UTR, or coding), predicted effect on protein sequence (e.g., synonymous, missense, nonsense, splice-site) and deleteriousness (LOW, MODERATE, HIGH).

To examine temporal changes in the frequency of recessive lethal alleles, we estimated annual allele frequencies from our complete dataset of hatched individuals (n=97). Annual allele frequencies were calculated from a single representative SNP at each locus – the missense variant in DLG1 and one of the splice-region variants in NEO1 – using all individuals living in a given year, with exact binomial confidence intervals computed to account for variation in annual sample size.

We explored potential structural variation (SV) on chromosome 10 by plotting sequencing depth for all samples. Per-sample sequencing depth in 100bp windows was calculated from aligned BAM files using *mosdepth* ^79^, and coverage profiles were visualized to identify patterns indicative of the SV type (e.g., inversion, duplication, deletion, etc).

To test whether carrier status at the DLG1 and NEO1 recessive lethal loci was associated with reduced reproductive output, we compared mean annual offspring production between carriers (individuals heterozygous or homozygous for the alternate allele) and non-carriers (homozygous reference) among hatched individuals. Annual offspring production was calculated by dividing total lifetime nestling production by lifespan in years. Differences in annual offspring production between carriers and non-carriers were tested separately for each locus using Wilcoxon rank-sum tests, given the non-normal distribution of offspring counts. For our complete dataset of breeding individuals, we observe no effect of recessive lethal carrier status on reproductive rates for DLG1 (n=80; p=0.61) and NEO1 (n=92, p=0.73) (**Fig. S12**). However, the likelihood that a carrier would breed with another carrier is very low when the allele is rare:for example, in the case of a recessive lethal with a 10% frequency, heterozygous carriers are expected to form breeding pairs only ∼2% of the time under random mating. Thus, to maximize the probability of detecting carrier effects, we restricted this analysis to individuals born after 2012 when the recessive lethal alleles were more abundant (**Fig. 3B**).

### Computational simulation model

We parameterized a simulation model using the non-Wright-Fisher (nonWF) model in SLiM 5 ^35,36^ to explore expected impacts of severe population bottleneck on deleterious genetic variation in ’Alalā. In the SLiM nonWF model, an individual’s absolute fitness governs its annual probabilities of survival and reproduction. Absolute fitness is determined by the cumulative effects of deleterious and beneficial genetic variants carried by an individual, with additional scaling applied to incorporate age-specific mortality.

We modeled deleterious mutations as nonsynonymous mutations in coding regions of a bird genome. The simulated genome consisted of 20,000 genes distributed across 40 chromosomes, with each gene spanning 2,000 bp, for a total of 40 Mb of coding sequence. Recombination was assumed to be absent within genes, to occur between genes at a rate of 1e-6 crossovers per site per generation, and to be effectively free among chromosomes. Deleterious mutations were introduced within genes at a rate of 3.5e-9 per site per generation, derived from an overall mutation rate of 5e-9 estimated for passerine birds ^37–39^ and assuming that 70% of coding mutations are nonsynonymous ^80^. To reduce computational overhead, synonymous (neutral) mutations were excluded from the simulations. Although some evidence points toward functional effects of synonymous variants ^81^, they typically have negligible impacts on fitness ^82^. Under these assumptions, the total deleterious mutation rate per diploid genome was U=0.28.

Selection (*s*) and dominance (*h*) coefficients for deleterious mutations followed the model described in Kyriazis et al ^40^. The vast majority of new mutations (99.5%) were drawn from a gamma distribution with mean selection coefficient *s* = -0.0131 and shape parameter 0.186, while the remaining 0.5% were assigned as fully recessive lethal mutations with *s* = -1.0 ^83^. Dominance coefficients were assigned as a function of selection strength, with *h* = 0.45 for weakly deleterious mutations (*s* > -0.001), *h* = 0.2 for moderately deleterious mutations (-0.001 ≥ s > -0.01), *h* = 0.05 for strongly deleterious mutations (-0.01 ≥ *s* > -0.1), and *h* = 0.0 for lethal and semi-lethal mutations (*s* ≤ −0.1). This parameterization yields a mean dominance coefficient of *h* = 0.28 for newly arising deleterious mutations, consistent with empirical estimates from human genetic data ^84^. Note that weakly deleterious (*s* > -0.001; *h*=0.45) comprise ∼49% of all new mutations in this model whereas strongly deleterious mutations (*s*<-0.01; *h*≤0.05) comprise ∼27%, enabling our simulation analysis to explore how the population bottleneck in ‘Alalā has impacted both on the mildly deleterious load as well as inbreeding depression due to strongly deleterious recessive mutations.

We set the life history parameters of our model based on information from the ’Alalā breeding program. Specifically, our model allows for overlapping generations where individuals are born each year, are reproductively active from age 2-18, and have a maximum lifespan of 25 years. These parameters are based on the observation that female clutch productivity declines close to zero after age 18 and nearly all captive individuals die by age 25^85^. Reproductive parameters were derived from our negative binomial GLM (see above), which estimated a mean annual number of nestlings of 0.66 for non-inbred individuals (F_ROH_=0) and dispersion parameter of 0.76. To model the impacts of inbreeding depression, we modeled annual reproduction for each reproductive individual as a function of their absolute fitness and then drew the number of nestlings from a successful reproductive event as 0.66*(maternal fitness)*(paternal fitness). For instance, if maternal and paternal fitnesses are both 0.95, the probability of reproducing in a given year would be ∼0.9 (0.95^2^) and the average number of nestlings produced per successful reproduction would be ∼0.6 (0.66*0.95^2^). Further, note that draws from a negative binomial with a mean of 0.6 and dispersion parameter of 0.76 yield 0 nestlings ∼63% of the time, 1 nestling ∼23% of the time, 2 nestlings ∼9% of the time, and >2 nestlings ∼5% of the time. Overall, these life history parameters yielded a generation time of 6 years for ’Alalā.

To model an ancestral ’Alalā population with a level of non-ROH heterozygosity that matched our empirical estimates (0.67 hets/kb), we determined the equilibrium effective population size (N_e_) that would yield this level of heterozygosity under a neutral model as N_e_ = 0.00067/(4*5e-9)=33,500. Note that this N_e_ estimate broadly agrees with our PSMC results, suggesting effective population sizes between ∼20,000-50,000 over the last 500,000 years prior to recent declines (**Fig. S7**). We converted this N_e_ estimate into a carrying capacity (K) by determining the ratio of N_e_:K during a neutral simulation with the above-described life history parameters (N_e_:K = 0.404). This ratio was used to compute the ancestral carrying capacity as K_ancestral_=33,500/0.404≈83,000. This ancestral carrying capacity determines the maximum potential population size throughout the initial burn-in period of our simulations, during which deleterious mutations were allowed to reach a near-equilibrium state over the course of 20,000 years (**Fig. S15**). To model an ancestral population with 3x greater level of non-ROH heterozygosity, we increased the per base pair deleterious mutation rate 3x from 3.5e-9 to 1.05e-8 (**Fig. S15**). For each of these mutation rate scenarios, we ran a single burn-in replicate and saved the simulation state for later use initiating replicates of different post-bottleneck simulation scenarios. This approach was implemented to lessen the computational burden of our simulations (each individual burn-in takes ∼5 days to complete) while still allowing for variability in the post-burn-in simulation outcomes.

Following an initial burn-in period, we imposed a two-stage population decline designed to reflect major phases of human impact in Hawaiʻi by introducing stochastic mortality. The first stage represented initial human arrival in Hawaiʻi consisting of an annual mortality probability of 0.17 that resulted in a gradual decline occurring over ∼550 years. This gradual decline was necessary to generate a baseline level of autozygosity such that simulated F_ROH_ closely matched empirical F_ROH_ following the population bottleneck (**Fig. 4**). The second stage of decline representing the arrival of Western colonists consisted of a much higher mortality probability of 0.23, occurring after the simulated population declined below 300 individuals. As the simulated wild population declined toward extinction, we founded a captive population by sampling individuals that exactly matched the sex and year of capture of the nine original founders (note that only eight of these founders still have descendants in the present-day population; see Hedrick et al. ^41^). These founders and their descendants were then allowed to survive and reproduce according to the stochastic dynamics of the model as the captive population increased to a carrying capacity of K=130, aiming to model a census size fluctuating around N=120 (note that N:K ratios are roughly determined by the absolute fitness of the simulated population). Additionally, note that the captive carrying capacity is already well beyond the current possible carrying capacity of K=88, as determined by the available number of aviaries. Because stochasticity often caused variation in recovery trajectories, we retained only simulation replicates in which the captive population size fell between 20 and 100 individuals in 2005 (note that the true population size in 2005 was ∼55 individuals).

Using this filtered model as a baseline, we evaluated three main scenarios: (1) a constant captive population carrying capacity (K = 130) with no releases into the wild; (2) a doubling of captive carrying capacity to K = 260 in 2030 with no wild releases; (3) a reintroduction scenario in which 10 individuals (<5 years in age) were released every year from 2026-2036 (100 individuals total) assuming an elevated annual mortality rate in the wild of 10%. To explore sensitivity to this assumed mortality rate increase as well as the number of reintroduced individuals, we also modeled two additional wild mortality rates (5% and 15%, with 100 individuals released) and two additional release quantities (50 or 150 individuals, assuming a 10% mortality rate increase). For each scenario, we ran 600 simulation replicates (of which ∼20% met acceptance criteria) and projected population dynamics from the present day through 2100. For each replicate and year, we recorded population size, mean F_ROH_ calculated from ROH >1 Mb, population fitness, the inbreeding load (measured as the diploid number of lethal equivalents ^40,42^), and the maximum frequency of recessive lethal mutations. Here, we focus on the maximum rather than average lethal frequency because the high-frequency lethal alleles we identify in our empirical analysis (**Fig. 3**) are likely to be outliers on the tail of the lethal frequency distribution and not representative of the full distribution of lethal alleles in ‘Alalā.

## References

1. Ceballos, G., Ehrlich, P. R. & Raven, P. H. Vertebrates on the brink as indicators of biological annihilation and the sixth mass extinction. Proc Natl Acad Sci U S A 117, 13596–13602 (2020).

2. Langhammer, P. F. et al. The positive impact of conservation action. Science 384, 453–458 (2024).

3. Wiedenfeld, D. A. et al. Conservation resource allocation, small population resiliency, and the fallacy of conservation triage. Conserv Biol 35, 1388–1395 (2021).

4. Kirkpatrick & Jarne. The effects of a bottleneck on inbreeding depression and the genetic load. Am. Nat. 155, 154 (2000).

5. López-Cortegano, E. & Charlesworth, B. The effect of a reduction in population size on mean fitness and inbreeding depression. bioRxiv (2026) doi:10.64898/2026.05.15.725556.

6. Jackson, H. A. et al. Genomic erosion in a demographically recovered bird species during conservation rescue. Conserv Biol 36, e13918 (2022).

7. Robinson, J. A. et al. Genome-wide diversity in the California condor tracks its prehistoric abundance and decline. Curr Biol 31, 2939–2946.e5 (2021).

8. Cavill, E. L. et al. When birds of a feather flock together: Severe genomic erosion and the implications for genetic rescue in an endangered island passerine. Evol Appl 17, e13739 (2024).

9. Fontsere, C. et al. Persistent Genomic Erosion in Whooping Cranes Despite Demographic Recovery. Mol Ecol e70088 (2025).

10. Dussex, N. et al. Population genomics of the critically endangered kākāpō. Cell Genom 1, 100002 (2021).

11. Tsujimoto, D. et al. Genetic purging in an island-endemic pigeon recovering from the brink of extinction. Commun Biol 8, 1051 (2025).

12. von Seth, J. et al. Genomic trajectories of a near-extinction event in the Chatham Island black robin. BMC Genomics 23, 747 (2022).

13. Femerling, G. et al. Genetic Load and Adaptive Potential of a Recovered Avian Species that Narrowly Avoided Extinction. Mol Biol Evol 40, (2023).

14. Kyriazis, C. C., Robinson, J. A. & Lohmueller, K. E. Long runs of homozygosity are reliable genomic markers of inbreeding depression. Trends Ecol Evol 40, 874–884 (2025).

15. Bertorelle, G. et al. Genetic load: genomic estimates and applications in non-model animals. Nat Rev Genet 23, 492–503 (2022).

16. Kyriazis, C. C. et al. Population genomics of recovery and extinction in Hawaiian honeycreepers. Curr Biol 35, 2697–2708.e4 (2025).

17. Fleischer, R. C., Campana, M. G. & James, H. F. Hawaiian songbird radiations. Curr Biol 32, R1070–R1072 (2022).

18. Blanchet, G. et al. Reduction of genetic diversity in ’Alalā (Hawaiian crow; Corvus hawaiiensis) between the late 1800s and the late 1900s. J Hered 115, 32–44 (2024).

19. Banko, W. E. & Banko, P. C. History of Endemic Hawaiian Birds. (1980).

20. Flanagan, A. M., Masuda, B., Grueber, C. E. & Sutton, J. T. Moving from trends to benchmarks by using regression tree analysis to find inbreeding thresholds in a critically endangered bird. Conserv Biol 35, 1278–1287 (2021).

21. Hoeck, P. E. A., Wolak, M. E., Switzer, R. A., Kuehler, C. M. & Lieberman, A. A. Effects of inbreeding and parental incubation on captive breeding success in Hawaiian crows. Biological Conservation 184, 357–364 (2015).

22. Heber, S. & Briskie, J. V. Population bottlenecks and increased hatching failure in endangered birds. Conserv Biol 24, 1674–1678 (2010).

23. Marshall, A. F., Balloux, F., Hemmings, N. & Brekke, P. Systematic review of avian hatching failure and implications for conservation. Biol Rev Camb Philos Soc 98, 807–832 (2023).

24. Larivière, D. et al. Scalable, accessible and reproducible reference genome assembly and evaluation in Galaxy. Nat Biotechnol 42, 367–370 (2024).

25. Sutton, J. T. et al. A High-Quality, Long-Read De Novo Genome Assembly to Aid Conservation of Hawaii’s Last Remaining Crow Species. Genes (Basel) 9, (2018).

26. Seppey, M., Manni, M. & Zdobnov, E. M. BUSCO: Assessing Genome Assembly and Annotation Completeness. Methods Mol Biol 1962, 227–245 (2019).

27. Robledo-Ruiz, D. A. et al. Chromosome-length genome assembly and linkage map of a critically endangered Australian bird: the helmeted honeyeater. Gigascience 11, (2022).

28. Formenti, G., et al. The Vertebrate Genomes Project Phase I: A global reference genome resource. bioRxivorg (2026) doi:10.64898/2026.06.24.732306.

29. Iizuka-Kogo, A. et al. Requirement of DLG1 for cardiovascular development and tissue elongation during cochlear, enteric, and skeletal development: possible role in convergent extension. PLoS One 10, e0123965 (2015).

30. Mahoney, Z. X. et al. Discs-large homolog 1 regulates smooth muscle orientation in the mouse ureter. Proc Natl Acad Sci U S A 103, 19872–19877 (2006).

31. Caruana, G. & Bernstein, A. Craniofacial dysmorphogenesis including cleft palate in mice with an insertional mutation in the discs large gene. Mol Cell Biol 21, 1475–1483 (2001).

32. Srinivasan, K., Strickland, P., Valdes, A., Shin, G. C. & Hinck, L. Netrin-1/neogenin interaction stabilizes multipotent progenitor cap cells during mammary gland morphogenesis. Dev Cell 4, 371–382 (2003).

33. Matsunaga, E. et al. RGM and its receptor neogenin regulate neuronal survival. Nat Cell Biol 6, 749–755 (2004).

34. Antin, P. B., Yatskievych, T. A., Davey, S. & Darnell, D. K. GEISHA: an evolving gene expression resource for the chicken embryo. Nucleic Acids Res 42, D933–7 (2014).

35. Haller, B. C. & Messer, P. W. SLiM 3: Forward Genetic Simulations Beyond the Wright-Fisher Model. Mol Biol Evol 36, 632–637 (2019).

36. Haller, B. C., Ralph, P. L. & Messer, P. W. SLiM 5: Eco-evolutionary simulations across multiple chromosomes and full genomes. Mol. Biol. Evol. 43, (2026).

37. Bergeron, L. A. et al. Evolution of the germline mutation rate across vertebrates. Nature 615, 285– 291 (2023).

38. Prentout, D. et al. Germline mutation rates and fine-scale recombination parameters in zebra finch. PLoS Genet 21, e1011661 (2025).

39. Smeds, L., Qvarnström, A. & Ellegren, H. Direct estimate of the rate of germline mutation in a bird. Genome Res 26, 1211–1218 (2016).

40. Kyriazis, C. C., Robinson, J. A. & Lohmueller, K. E. Using Computational Simulations to Model Deleterious Variation and Genetic Load in Natural Populations. Am Nat 202, 737–752 (2023).

41. Hedrick, P. W., Hoeck, P. E. A., Fleischer, R. C., Farabaugh, S. & Masuda, B. M. The influence of captive breeding management on founder representation and inbreeding in the ‘Alalā, the Hawaiian crow. Conserv. Genet. 17, 369–378 (2016).

42. Morton, N. E., Crow, J. F. & Muller, H. J. An Estimate of the Mutational Damage in Man from Data on Consanguineous Marriages. Proc Natl Acad Sci U S A 42, 855–863 (1956).

43. Nietlisbach, P., Muff, S., Reid, J. M., Whitlock, M. C. & Keller, L. F. Nonequivalent lethal equivalents: Models and inbreeding metrics for unbiased estimation of inbreeding load. Evol Appl 12, 266–279 (2019).

44. Greggor, A. L. et al. Balancing evidence and reducing uncertainty in the evaluation of reintroduction outcomes in ‘alalā, the Hawaiian crow. Global Ecology and Conservation 62, e03673 (2025).

45. Grossen, C., Guillaume, F., Keller, L. F. & Croll, D. Purging of highly deleterious mutations through severe bottlenecks in Alpine ibex. Nat Commun 11, 1001 (2020).

46. Khan, A. et al. Genomic evidence for inbreeding depression and purging of deleterious genetic variation in Indian tigers. Proc Natl Acad Sci U S A 118, (2021).

47. Dussex, N., Morales, H. E., Grossen, C., Dalén, L. & van Oosterhout, C. Purging and accumulation of genetic load in conservation. Trends Ecol Evol 38, 961–969 (2023).

48. Lynch, M., Conery, J. & Burger, R. Mutation accumulation and the extinction of small populations. Am. Nat. 146, 489–518 (1995).

49. Niehaus, J. et al. Evolutionary dynamics of a lethal recessive allele in reintroduced fragmented lynx populations. bioRxiv (2025) doi:10.1101/2025.11.06.686959.

50. Tian, D., et al. Long-term small effective population size, inbreeding, and a recessive lethal haplotype drive premature death in the endangered Devils Hole pupfish (Cyprinodon diabolis). bioRxivorg (2026) doi:10.64898/2026.06.06.730634.

51. Kennedy, E. S., Grueber, C. E., Duncan, R. P. & Jamieson, I. G. Severe inbreeding depression and no evidence of purging in an extremely inbred wild species--the Chatham Island black robin. Evolution 68, 987–995 (2014).

52. Marion, S. B. & Noor, M. A. F. Interrogating the Roles of Mutation-Selection Balance, Heterozygote Advantage, and Linked Selection in Maintaining Recessive Lethal Variation in Natural Populations. Annu Rev Anim Biosci 11, 77–91 (2023).

53. Stoffel, M. A., Johnston, S. E., Pilkington, J. G. & Pemberton, J. M. Purifying and balancing selection on embryonic semi-lethal haplotypes in a wild mammal. Evol Lett 8, 222–230 (2024).

54. Ober, C., Hyslop, T. & Hauck, W. W. Inbreeding effects on fertility in humans: evidence for reproductive compensation. Am J Hum Genet 64, 225–231 (1999).

55. Foster, Y. et al. Genomic Architecture of Inbreeding Depression Associated With Hatching Failure in an Endangered Parrot. Mol Ecol 35, e70252 (2026).

56. Robledo-Ruiz, D. A. et al. Genetic load proxies do not predict fitness better than inbreeding does in a wild population. bioRxiv (2025) doi:10.1101/2025.07.07.663445.

57. Robinson, J. A. et al. The critically endangered vaquita is not doomed to extinction by inbreeding depression. Science 376, 635–639 (2022).

58. Kardos, M., Keller, L. F. & Funk, W. C. What Can Genome Sequence Data Reveal About Population Viability? Mol Ecol 34, e17608 (2025).

59. Kyriazis, C. C. et al. Models based on best-available information support a low inbreeding load and potential for recovery in the vaquita. Heredity (Edinb) 130, 183–187 (2023).

60. Formenti, G. et al. Complete vertebrate mitogenomes reveal widespread repeats and gene duplications. Genome Biol. 22, 120 (2021).

61. Weissensteiner, M. H. et al. Combination of short-read, long-read, and optical mapping assemblies reveals large-scale tandem repeat arrays with population genetic implications. Genome Res 27, 697–708 (2017).

62. Dussex, N. et al. A genome-wide investigation of adaptive signatures in protein-coding genes related to tool behaviour in New Caledonian and Hawaiian crows. Mol Ecol 30, 973–986 (2021).

63. Weissensteiner, M. H. et al. Discovery and population genomics of structural variation in a songbird genus. Nat Commun 11, 3403 (2020).

64. Harris, R. S. Improved Pairwise Alignment of Genomic DNA. (2007).

65. Krzywinski, M. et al. Circos: an information aesthetic for comparative genomics. Genome Res 19, 1639–1645 (2009).

66. Van der Auwera, G. A. et al. From FastQ data to high confidence variant calls: the Genome Analysis Toolkit best practices pipeline. Curr Protoc Bioinformatics 43, 11.10.1–11.10.33 (2013).

67. Martin, M. Cutadapt removes adapter sequences from high-throughput sequencing reads. EMBnet J. (2011) doi:10.14806/ej.17.1.200.

68. Li, H. Aligning sequence reads, clone sequences and assemblycontigs with BWA-MEM. arXiv (2013) doi:10.48550/arXiv.1303.3997.

69. Tarasov, A., Vilella, A. J., Cuppen, E., Nijman, I. J. & Prins, P. Sambamba: fast processing of NGS alignment formats. Bioinformatics 31, 2032–2034 (2015).

70. Benson, G. Tandem repeats finder: a program to analyze DNA sequences. Nucleic Acids Res 27, 573–580 (1999).

71. Morgulis, A., Gertz, E. M., Schäffer, A. A. & Agarwala, R. WindowMasker: window-based masker for sequenced genomes. Bioinformatics 22, 134–141 (2006).

72. R Core Team. R: A Language and Environment for Statistical Computing. Preprint at https://www.R-project.org/ (2025).

73. Korunes, K. L. & Samuk, K. pixy: Unbiased estimation of nucleotide diversity and divergence in the presence of missing data. Mol Ecol Resour 21, 1359–1368 (2021).

74. Narasimhan, V. et al. BCFtools/RoH: a hidden Markov model approach for detecting autozygosity from next-generation sequencing data. Bioinformatics 32, 1749–1751 (2016).

75. Lacy, R. C., Ballou, J. D. & Pollak, J. P. PMx: software package for demographic and genetic analysis and management of pedigreed populations. Methods Ecol. Evol. 3, 433–437 (2012).

76. Li, H. & Durbin, R. Inference of human population history from individual whole-genome sequences. Nature 475, 493–496 (2011).

77. Hilgers, L. et al. Avoidable false PSMC population size peaks occur across numerous studies. Curr Biol 35, 927–930.e3 (2025).

78. Cingolani, P. et al. A program for annotating and predicting the effects of single nucleotide polymorphisms, SnpEff: SNPs in the genome of Drosophila melanogaster strain w1118; iso-2; iso-3. Fly (Austin) 6, 80–92 (2012).

79. Pedersen, B. S. & Quinlan, A. R. Mosdepth: quick coverage calculation for genomes and exomes. Bioinformatics 34, 867–868 (2018).

80. Huber, C. D., Kim, B. Y., Marsden, C. D. & Lohmueller, K. E. Determining the factors driving selective effects of new nonsynonymous mutations. Proc Natl Acad Sci U S A 114, 4465–4470 (2017).

81. Radrizzani, S., Kudla, G., Izsvák, Z. & Hurst, L. D. Selection on synonymous sites: the unwanted transcript hypothesis. Nat. Rev. Genet. 25, 431–448 (2024).

82. Dhindsa, R. S. et al. A minimal role for synonymous variation in human disease. Am. J. Hum. Genet. 109, 2105–2109 (2022).

83. Wade, E. E., Kyriazis, C. C., Cavassim, M. I. A. & Lohmueller, K. E. Quantifying the fraction of new mutations that are recessive lethal. Evolution 77, 1539–1549 (2023).

84. Kyriazis, C. C. & Lohmueller, K. E. Constraining models of dominance for nonsynonymous mutations in the human genome. PLoS Genet 20, e1011198 (2024).

85. Lisa P. Barrett, Alison M. Flanagan, Bryce Masuda, Ronald R. Swaisgood. The influence of pair duration on reproductive success in the monogamous ‘Alalā (Hawaiian crow). Frontiers in Conservation Science 24, (2024).

