## Supplementary Information for "Elevated recessive lethal frequencies drive hatching failure following near extinction in ‘Alalā, the Hawaiian crow"

**Table S1: Summary of sequenced individuals.** AL samples are hatched individuals and K samples are unhatched embryos. DLG1 and NEO1 genotypes encoded as 0 for homozygous reference, 1 for heterozygous, 2 for homozygous alternate, and blank for missing data.

| # | Sample | SRA Accession | Hatch Year | Depth | $F_{ROH}$ | DLG1 | NEO1 |
|---|---|---|---|---|---|---|---|
| 1 | AL032 | SAMN61868873 | 1989 | 18.94 | 0.47 | 0 | 0 |
| 2 | AL035 | SAMN61868874 | 1992 | 21.17 | 0.24 | 0 | 0 |
| 3 | AL075 | SAMN61868875 | 1997 | 20.8 | 0.18 | 1 | 1 |
| 4 | AL082 | SAMN61868876 | 1997 | 22.09 | 0.38 | 0 | 1 |
| 5 | AL086 | SAMN61868877 | 1998 | 14.31 | 0.27 | 0 | 0 |
| 6 | AL088 | SAMN61868878 | 1999 | 17.44 | 0.37 | 0 | 1 |
| 7 | AL091 | SAMN61868879 | 2000 | 15.07 | 0.3 | 0 | 0 |
| 8 | AL092 | SAMN61868880 | 2000 | 16.18 | 0.26 | 0 | 0 |
| 9 | AL095 | SAMN61868881 | 2001 | 24.71 | 0.15 | 1 | 0 |
| 10 | AL097 | SAMN61868882 | 2001 | 24.74 | 0.29 | 1 | 0 |
| 11 | AL099 | SAMN61868883 | 2001 | 22.01 | 0.24 | 1 | 1 |
| 12 | AL101 | SAMN61868884 | 2001 | 18.72 | 0.31 | 1 | 0 |
| 13 | AL102 | SAMN61868885 | 2002 | 1.032 | 0.36 | 0 | 0 |
| 14 | AL103 | SAMN61868886 | 2002 | 3.091 | 0.26 | 1 | 0 |
| 15 | AL114 | SAMN61868887 | 2004 | 23.66 | 0.43 | 0 | 0 |
| 16 | AL119 | SAMN61868888 | 2004 | 24.06 | 0.18 | 0 | 0 |
| 17 | AL120 | SAMN61868889 | 2004 | 13.61 | 0.33 | 0 | 0 |
| 18 | AL121 | SAMN61868890 | 2004 | 16.26 | 0.37 | 0 | 1 |

| | | | | | | | |
|---|---|---|---|---|---|---|---|
| 19 | AL124 | SAMN61868891 | 2004 | 23.62 | 0.36 | 0 | 1 |
| 20 | AL125 | SAMN61868892 | 2004 | 3.179 | 0.17 | 0 | 0 |
| 21 | AL127 | SAMN61868893 | 2005 | 2.847 | 0.32 | 1 | 0 |
| 22 | AL129 | SAMN61868894 | 2005 | 25.06 | 0.29 | 1 | 1 |
| 23 | AL133 | SAMN61868895 | 2005 | 2.574 | 0.33 | 1 | 0 |
| 24 | AL134 | SAMN61868896 | 2006 | 15.83 | 0.39 | | |
| 25 | AL137 | SAMN61868897 | 2006 | 29.1 | 0.2 | | 0 |
| 26 | AL138 | SAMN61868898 | 2007 | 16.78 | 0.21 | 1 | 1 |
| 27 | AL141 | SAMN61868899 | 2007 | 14.3 | 0.24 | 1 | 0 |
| 28 | AL142 | SAMN61868900 | 2007 | 18.61 | 0.18 | 1 | 0 |
| 29 | AL143 | SAMN61868901 | 2007 | 21.56 | 0.26 | 1 | 0 |
| 30 | AL146 | SAMN61868902 | 2008 | 18.93 | 0.27 | 1 | 1 |
| 31 | AL149 | SAMN61868903 | 2008 | 15.03 | 0.28 | | 0 |
| 32 | AL153 | SAMN61868904 | 2009 | 22.9 | 0.31 | 0 | |
| 33 | AL155 | SAMN61868905 | 2009 | 17.79 | 0.31 | 1 | 0 |
| 34 | AL156 | SAMN61868906 | 2009 | 2.629 | 0.35 | | 0 |
| 35 | AL159 | SAMN61868907 | 2010 | 15.82 | 0.27 | 0 | 1 |
| 36 | AL160 | SAMN61868908 | 2010 | 1.523 | 0.27 | 1 | 0 |
| 37 | AL162 | SAMN61868909 | 2010 | 16.11 | 0.23 | 1 | 1 |
| 38 | AL166 | SAMN61868910 | 2010 | 16.54 | 0.23 | 1 | 0 |
| 39 | AL170 | SAMN61868911 | 2010 | 16.94 | 0.37 | 0 | 0 |

| | | | | | | | |
|---|---|---|---|---|---|---|---|
| 40 | AL173 | SAMN61868912 | 2011 | 21.67 | 0.33 | 0 | 0 |
| 41 | AL174 | SAMN61868913 | 2011 | 14.78 | 0.24 | 0 | 0 |
| 42 | AL175 | SAMN61868914 | 2011 | 15.65 | 0.34 | 0 | 1 |
| 43 | AL176 | SAMN61868915 | 2011 | 14.66 | 0.31 | 0 | 0 |
| 44 | AL177 | SAMN61868916 | 2011 | 18.21 | 0.27 | 0 | 0 |
| 45 | AL178 | SAMN61868917 | 2011 | 15.25 | 0.25 | | 1 |
| 46 | AL180 | SAMN61868918 | 2011 | 16.14 | 0.29 | 0 | 0 |
| 47 | AL181 | SAMN61868919 | 2011 | 14.57 | 0.33 | | |
| 48 | AL182 | SAMN61868920 | 2011 | 21.39 | 0.3 | 0 | 0 |
| 49 | AL184 | SAMN61868921 | 2011 | 14.11 | 0.35 | 0 | 1 |
| 50 | AL185 | SAMN61868922 | 2011 | 2.866 | 0.29 | 0 | 0 |
| 51 | AL187 | SAMN61868923 | 2011 | 15.12 | 0.27 | 1 | 0 |
| 52 | AL189 | SAMN61868924 | 2011 | 23.87 | 0.31 | 1 | 1 |
| 53 | AL193 | SAMN61868925 | 2012 | 16.56 | 0.3 | 0 | 0 |
| 54 | AL196 | SAMN61868926 | 2012 | 21.87 | 0.27 | 1 | 0 |
| 55 | AL197 | SAMN61868927 | 2012 | 16.22 | 0.22 | 1 | 1 |
| 56 | AL199 | SAMN61868928 | 2012 | 21.41 | 0.32 | 0 | 1 |
| 57 | AL200 | SAMN61868929 | 2012 | 6.854 | 0.36 | 1 | 0 |
| 58 | AL201 | SAMN61868930 | 2012 | 17.39 | 0.3 | 1 | 1 |
| 59 | AL204 | SAMN61868931 | 2012 | 22.19 | 0.39 | 1 | 1 |
| 60 | AL207 | SAMN61868932 | 2012 | 3.482 | 0.25 | 1 | 1 |

| | | | | | | | |
|---|---|---|---|---|---|---|---|
| 61 | AL208 | SAMN61868933 | 2012 | 3.771 | 0.33 | | 0 |
| 62 | AL209 | SAMN61868934 | 2013 | 17.02 | 0.28 | 0 | 0 |
| 63 | AL210 | SAMN61868935 | 2013 | 13.61 | 0.32 | 1 | 0 |
| 64 | AL213 | SAMN61868936 | 2013 | 15.07 | 0.28 | 0 | 0 |
| 65 | AL215 | SAMN61868937 | 2013 | 20.69 | 0.36 | 0 | 0 |
| 66 | AL216 | SAMN61868938 | 2013 | 2.455 | 0.48 | 0 | 0 |
| 67 | AL220 | SAMN61868939 | 2014 | 3.441 | 0.34 | 1 | 1 |
| 68 | AL223 | SAMN61868940 | 2014 | 28.45 | 0.27 | | 0 |
| 69 | AL231 | SAMN61868941 | 2015 | 17.87 | 0.36 | 1 | 0 |
| 70 | AL238 | SAMN61868942 | 2015 | 17.64 | 0.29 | 0 | 1 |
| 71 | AL241 | SAMN61868943 | 2016 | 22.43 | 0.32 | | |
| 72 | AL242 | SAMN61868944 | 2016 | 2.788 | 0.35 | | 0 |
| 73 | AL243 | SAMN61868945 | 2016 | 18.04 | 0.39 | 1 | 0 |
| 74 | AL244 | SAMN61868946 | 2016 | 17.86 | 0.32 | 1 | 1 |
| 75 | AL246 | SAMN61868947 | 2016 | 3.127 | 0.35 | 0 | 0 |
| 76 | AL254 | SAMN61868948 | 2016 | 36.49 | 0.34 | 0 | 1 |
| 77 | AL255 | SAMN61868949 | 2016 | 27.3 | 0.43 | | 1 |
| 78 | AL256 | SAMN61868950 | 2016 | 27.14 | 0.33 | | 1 |
| 79 | AL263 | SAMN61868951 | 2016 | 21.56 | 0.28 | 0 | 1 |
| 80 | AL266 | SAMN61868952 | 2016 | 16.42 | 0.36 | 0 | 0 |
| 81 | AL269 | SAMN61868953 | 2016 | 14.43 | 0.21 | 1 | 0 |

| | | | | | | | |
|---|---|---|---|---|---|---|---|
| 82 | AL272 | SAMN61868954 | 2017 | 4.81 | 0.27 | 0 | 0 |
| 83 | AL273 | SAMN61868955 | 2017 | 10.52 | 0.29 | | 0 |
| 84 | AL274 | SAMN61868956 | 2017 | 17.18 | 0.34 | 0 | 0 |
| 85 | AL276 | SAMN61868957 | 2017 | 16.84 | 0.36 | 1 | 1 |
| 86 | AL278 | SAMN61868958 | 2017 | 24.79 | 0.26 | | 1 |
| 87 | AL279 | SAMN61868959 | 2017 | 19.56 | 0.4 | 1 | 1 |
| 88 | AL283 | SAMN61868960 | 2017 | 13.76 | 0.35 | 0 | 1 |
| 89 | AL284 | SAMN61868961 | 2017 | 18.69 | 0.35 | 0 | 0 |
| 90 | AL288 | SAMN61868962 | 2017 | 19.02 | 0.3 | 1 | 0 |
| 91 | AL292 | SAMN61868963 | 2017 | 23.87 | 0.4 | 1 | 1 |
| 92 | AL300 | SAMN61868964 | 2017 | 19.03 | 0.39 | 0 | 0 |
| 93 | AL303 | SAMN61868965 | 2017 | 19.16 | 0.37 | | 1 |
| 94 | AL304 | SAMN61868966 | 2017 | 24.59 | 0.32 | | 0 |
| 95 | AL307 | SAMN61868967 | 2018 | 20.28 | 0.34 | 0 | 0 |
| 96 | AL309 | SAMN61868968 | 2018 | 19.18 | 0.34 | 1 | 0 |
| 97 | AL313 | SAMN61868969 | 2018 | 19.62 | 0.36 | 1 | 0 |
| 98 | K1423 | SAMN61868970 | 2014 | 13.59 | 0.27 | 1 | 1 |
| 99 | K1461 | SAMN61868971 | 2014 | 19.48 | 0.31 | 0 | 0 |
| 100 | K1542 | SAMN61868972 | 2015 | 17.35 | 0.32 | 2 | 1 |
| 101 | K1544 | SAMN61868973 | 2015 | 13.62 | 0.33 | 1 | 0 |
| 102 | K1567 | SAMN61868974 | 2015 | 23.13 | 0.27 | 2 | 0 |

| | | | | | | | |
|---|---|---|---|---|---|---|---|
| 103 | K1579 | SAMN61868975 | 2015 | 43.88 | 0.25 | 1 | 0 |
| 104 | K1580 | SAMN61868976 | 2015 | 15.35 | 0.38 | 0 | 0 |
| 105 | K1583 | SAMN61868977 | 2015 | 18.41 | 0.33 | 1 | 2 |
| 106 | K1603 | SAMN61868978 | 2016 | 16.22 | 0.32 | 0 | 0 |
| 107 | K1606 | SAMN61868979 | 2016 | 26.31 | 0.3 | 0 | 2 |
| 108 | K1608 | SAMN61868980 | 2016 | 24.57 | 0.26 | 0 | 0 |
| 109 | K16100 | SAMN61868981 | 2016 | 19.12 | 0.37 | 1 | 1 |
| 110 | K1614 | SAMN61868982 | 2016 | 16.23 | 0.26 | | 2 |
| 111 | K1617 | SAMN61868983 | 2016 | 24.56 | 0.33 | 0 | 0 |
| 112 | K1618 | SAMN61868984 | 2016 | 22.02 | 0.39 | 0 | 2 |
| 113 | K1623 | SAMN61868985 | 2016 | 3.664 | 0.34 | 0 | |
| 114 | K1627 | SAMN61868986 | 2016 | 5.889 | 0.36 | | 2 |
| 115 | K1632 | SAMN61868987 | 2016 | 18.77 | 0.37 | 2 | 2 |
| 116 | K1640 | SAMN61868988 | 2016 | 26.13 | 0.42 | 0 | 0 |
| 117 | K1650 | SAMN61868989 | 2016 | 18.14 | 0.27 | 0 | 0 |
| 118 | K1660 | SAMN61868990 | 2016 | 10.35 | 0.38 | 0 | 1 |
| 119 | K1663 | SAMN61868991 | 2016 | 12.39 | 0.49 | 0 | 2 |
| 120 | K1665 | SAMN61868992 | 2016 | 20.05 | 0.26 | 1 | 0 |
| 121 | K1669 | SAMN61868993 | 2016 | 29.44 | 0.3 | 0 | 0 |
| 122 | K1672 | SAMN61868994 | 2016 | 30.68 | 0.34 | 0 | 0 |
| 123 | K1680 | SAMN61868995 | 2016 | 18.25 | 0.31 | 0 | 0 |

| | | | | | | | |
|---|---|---|---|---|---|---|---|
| 124 | K1684 | SAMN61868996 | 2016 | 29.14 | 0.27 | 1 | 0 |
| 125 | K1693 | SAMN61868997 | 2016 | 14.38 | 0.29 | 0 | 0 |
| 126 | K1696 | SAMN61868998 | 2016 | 17.08 | 0.26 | 0 | 2 |
| 127 | K17007 | SAMN61868999 | 2017 | 16.6 | 0.33 | 0 | 0 |
| 128 | K17008 | SAMN61869000 | 2017 | 2.673 | 0.42 | | |
| 129 | K17010 | SAMN61869001 | 2017 | 24.89 | 0.36 | 0 | 0 |
| 130 | K17013 | SAMN61869002 | 2017 | 9.852 | 0.31 | | 0 |
| 131 | K17014 | SAMN61869003 | 2017 | 15.69 | 0.32 | 1 | 0 |
| 132 | K17025 | SAMN61869004 | 2017 | 18.59 | 0.29 | 1 | 0 |
| 133 | K17028 | SAMN61869005 | 2017 | 16.66 | 0.29 | 0 | 2 |
| 134 | K17030 | SAMN61869006 | 2017 | 16.31 | 0.36 | 0 | 0 |
| 135 | K17032 | SAMN61869007 | 2017 | 41.34 | 0.42 | 0 | 2 |
| 136 | K17033 | SAMN61869008 | 2017 | 32.02 | 0.38 | 2 | 2 |
| 137 | K17034 | SAMN61869009 | 2017 | 15.05 | 0.3 | 0 | 0 |
| 138 | K17035 | SAMN61869010 | 2017 | 12.75 | 0.23 | 1 | 0 |
| 139 | K17037 | SAMN61869011 | 2017 | 16.21 | 0.33 | 0 | 1 |
| 140 | K17040 | SAMN61869012 | 2017 | 22.04 | 0.38 | 1 | 2 |
| 141 | K17042 | SAMN61869013 | 2017 | 22.92 | 0.38 | 1 | 0 |
| 142 | K17044 | SAMN61869014 | 2017 | 23.17 | 0.26 | 1 | 0 |
| 143 | K17045 | SAMN61869015 | 2017 | 87.19 | 0.4 | 1 | 0 |
| 144 | K17048 | SAMN61869016 | 2017 | 15.49 | 0.42 | 1 | 0 |

| | | | | | | | |
|---|---|---|---|---|---|---|---|
| 145 | K17052 | SAMN61869017 | 2017 | 33.54 | 0.35 | 0 | 0 |
| 146 | K17053 | SAMN61869018 | 2017 | 17.9 | 0.42 | 1 | 1 |
| 147 | K17056 | SAMN61869019 | 2017 | 24.55 | 0.37 | 1 | 1 |
| 148 | K17062 | SAMN61869020 | 2017 | 20.28 | 0.34 | 1 | 1 |
| 149 | K17064 | SAMN61869021 | 2017 | 16.4 | 0.23 | 1 | 1 |
| 150 | K17073 | SAMN61869022 | 2017 | 17.82 | 0.31 | 1 | 2 |
| 151 | K17077 | SAMN61869023 | 2017 | 17.61 | 0.29 | 0 | 0 |
| 152 | K17078 | SAMN61869024 | 2017 | 18.48 | 0.36 | 0 | 0 |
| 153 | K17081 | SAMN61869025 | 2017 | 21.04 | 0.31 | 0 | 0 |
| 154 | K17083 | SAMN61869026 | 2017 | 19.73 | 0.39 | 2 | 0 |
| 155 | K17087 | SAMN61869027 | 2017 | 25.5 | 0.37 | 0 | 0 |
| 156 | K17091 | SAMN61869028 | 2017 | 18.23 | 0.39 | 2 | 1 |
| 157 | K17094 | SAMN61869029 | 2017 | 22.1 | 0.39 | 0 | 0 |
| 158 | K17098 | SAMN61869030 | 2017 | 18.07 | 0.26 | 1 | 1 |
| 159 | K17099 | SAMN61869031 | 2017 | 17.57 | 0.35 | 0 | 2 |
| 160 | K17101 | SAMN61869032 | 2017 | 26.49 | 0.3 | 2 | 0 |
| 161 | K17102 | SAMN61869033 | 2017 | 20.65 | 0.26 | 1 | 1 |
| 162 | K17103 | SAMN61869034 | 2017 | 31.7 | 0.36 | | 1 |
| 163 | K17104 | SAMN61869035 | 2017 | 17.98 | 0.31 | 0 | 0 |
| 164 | K17106 | SAMN61869036 | 2017 | 21.95 | 0.34 | 2 | 0 |
| 165 | K18010 | SAMN61869037 | 2018 | 15.37 | 0.37 | 1 | 1 |

| | | | | | | | |
|---|---|---|---|---|---|---|---|
| 166 | K18017 | SAMN61869038 | 2018 | 24.44 | 0.34 | 1 | 1 |
| 167 | K18019 | SAMN61869039 | 2018 | 22.05 | 0.25 | 0 | 1 |
| 168 | K18022 | SAMN61869040 | 2018 | 17.5 | 0.4 | 0 | 2 |
| 169 | K18034 | SAMN61869041 | 2018 | 11.02 | 0.35 | | 1 |
| 170 | K18078 | SAMN61869042 | 2018 | 20.12 | 0.34 | 1 | 1 |
| 171 | K18083 | SAMN61869043 | 2018 | 17.44 | 0.31 | | 0 |
| 172 | K18084 | SAMN61869044 | 2018 | 16.59 | 0.34 | 0 | 1 |
| 173 | K18091 | SAMN61869045 | 2018 | 16.4 | 0.34 | 1 | 0 |
| 174 | K19018 | SAMN61869046 | 2019 | 17.4 | 0.32 | | 1 |
| 175 | K19019 | SAMN61869047 | 2019 | 15.83 | 0.33 | 1 | 1 |

**Table S2: Summary of Cox proportional hazards models relating $F_{ROH}$ (for ROH thresholds of 100kb, 1Mb, and 10Mb) to risk of death while controlling for hatch year as a frailty term.** Note that in both models hatch year is highly correlated with survival ($p<1e-5$).

| variable | n | coefficient | S.E. | p |
|---|---|---|---|---|
| FROH_100kb | 97 | 0.450 | 2.254 | 0.842 |
| FROH_1Mb | 97 | -0.0634 | 2.040 | 0.975 |
| FROH_10Mb | 97 | -1.828 | 2.474 | 0.46 |
| FROH_1Mb_highcov | 81 | 0.658 | 2.163 | 0.761 |

**Table S3: Relationship between $F_{ROH}$ and reproductive success in mixed effects negative binomial models**. In addition to the fixed effect (Variable), all models also included the adult's hatch year as a random effect and its longevity as an offset. FROH_100kb, FROH_1Mb, FROH_1Mb_highcov, and FROH_10Mb are estimated from empirical data (see Figs. 2E, S3E, S4E, and S6E) and nonROH_0.67 and nonROH_2.0 are generated from simulated data (see Fig. 5).

| Variable | n | Coefficient | Intercept | S.E. | z | p |
|---|---|---|---|---|---|---|
| FROH_100kb | 97 | -3.401 | 0.0571 | 2.079 | -1.636 | 0.102 |
| FROH_1Mb | 97 | -2.662 | -0.418 | 1.955 | -1.361 | 0.173 |
| FROH_1Mb_highcov | 81 | -1.302 | -0.881 | 2.428 | -0.536 | 0.592 |
| FROH_10Mb | 97 | -1.808 | -1.059 | 2.254 | -0.802 | 0.422 |
| nonROH_0.67 | 123 | -1.238 | -0.9428 | 0.892 | -1.387 | 0.165 |
| nonROH_2.0 | 111 | -5.689 | -0.7094 | 1.745 | -3.26 | 0.00111 |

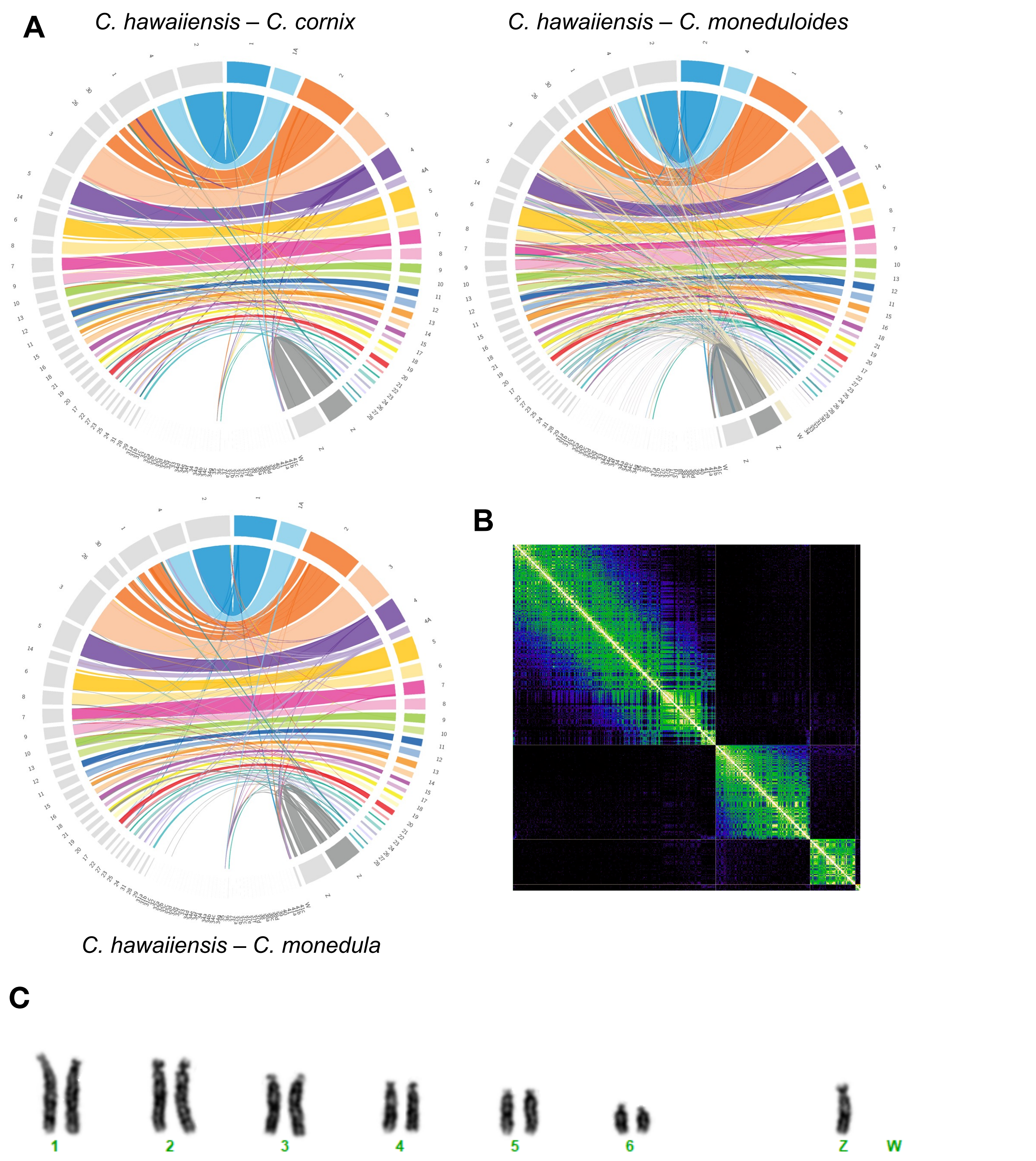


**Figure S1: Visualization of genome assembly synteny and macrochromosome karyotypes.** (A) Whole genome alignments for ʻAlalā and three different corvid species. Colors designate different chromosomes, with ʻAlalā shown on the left in each figure. Note fragmentation of Chromosome 2 in the ʻAlalā. (B) Hi-C contact frequency plot further supporting the fragmentation of Chromosome 2 in 'Alalā. The three distinct interaction blocks along the diagonal, separated by regions of near-zero contact frequency, indicate that the three fragments behave as independent chromosomes rather than a single continuous chromosome. (C) Giemsa-stained metaphase chromosomes of a female (ZW) ʻAlalā showing the six largest autosomal pairs and the ZW sex chromosomes. Note that chromosome numbering in (A) follows corvid synteny-based naming conventions, while numbering in (C) reflects descending size order as is standard in cytogenetic analyses.

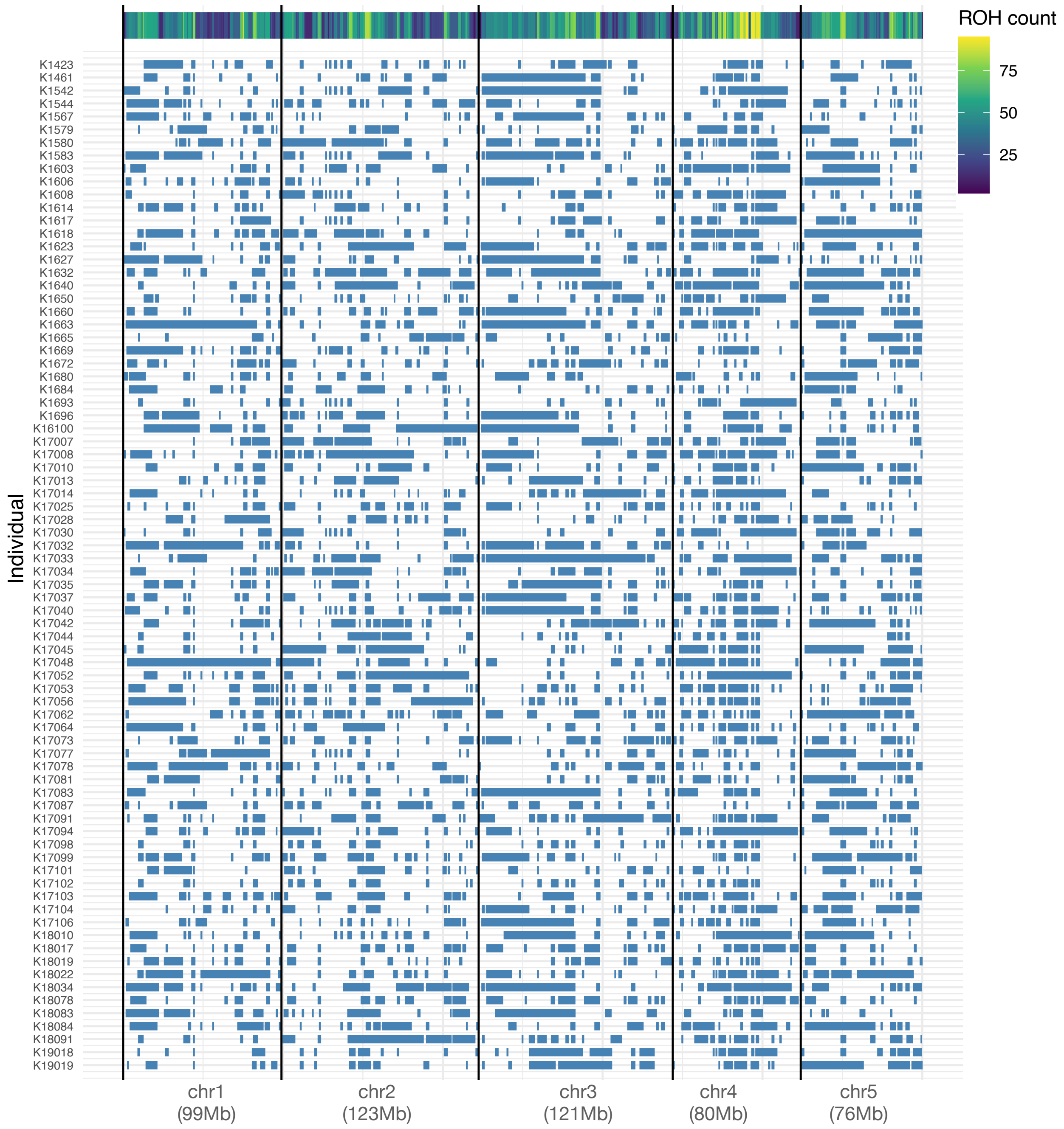


**Figure S2:** Runs of homozygosity visualized across the first five chromosomes for all 78 ‘Alalā embryos that failed to hatch. Heatmap of ROH abundance shown on the top.

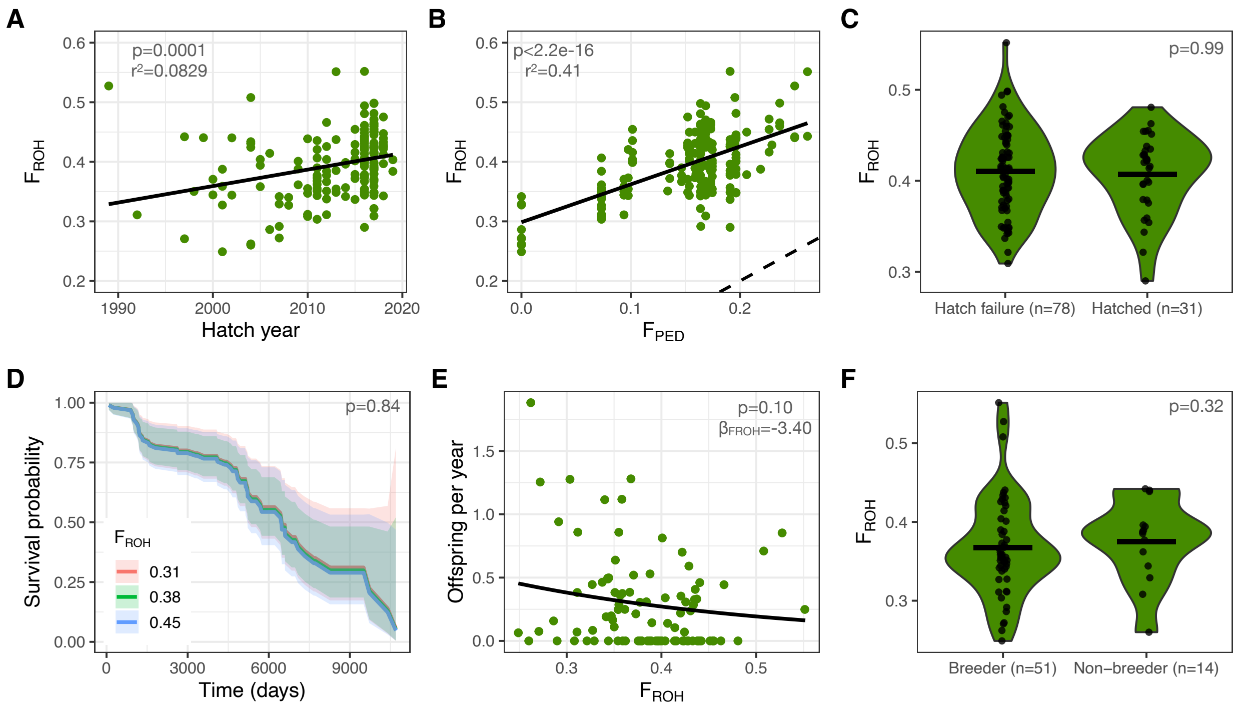


**Figure S3:** Patterns of inbreeding and inbreeding depression in the 'Alalā conservation breeding program when assuming an ROH threshold of 100kb. (A) Levels of inbreeding (measured by $F_{ROH}$ for ROH > 100kb) minimally increase over time and are elevated even in individuals hatched in the 1990s. (B) $F_{ROH}$ is correlated with, though much greater than, $F_{ped}$. Dashed line shows 1:1 relationship. (C) Levels of $F_{ROH}$ do not differ between 78 embryos that failed to hatch and 31 hatched individuals born during the same time period. (D) No impact of $F_{ROH}$ on adult survival in a Cox proportional hazards model (p=0.84). See Table S2 for full model output. (E) $F_{ROH}$ is negatively correlated with annual reproductive output, though this relationship is not statistically significant. See Table S3 for full model output. (F) No differences in $F_{ROH}$ between 51 individuals that produced viable offspring at any point in their life versus 14 individuals that never successfully reproduced. Note that horizontal black lines in panels C and F denote the means of each distribution.

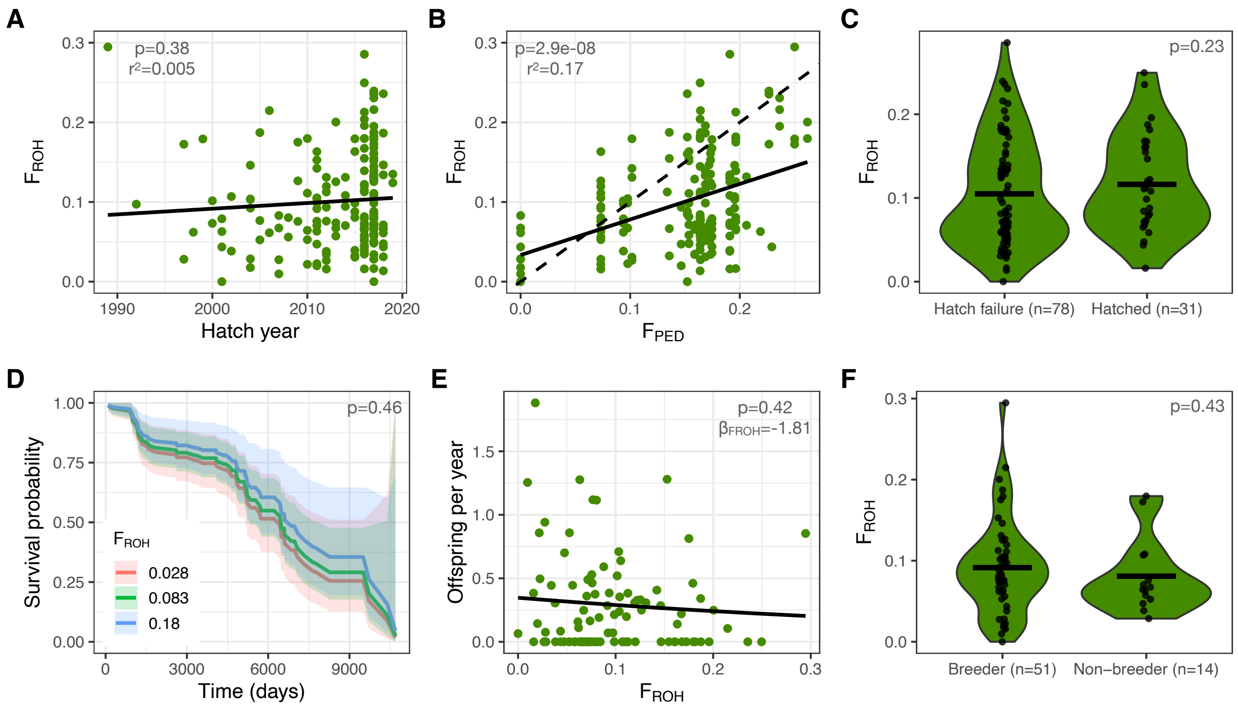


**Figure S4:** Patterns of inbreeding and inbreeding depression in the 'Alalā conservation breeding program when assuming an ROH threshold of 10Mb. (A) Levels of inbreeding (measured by $F_{ROH}$ for ROH > 10Mb) minimally increase over time and are elevated even in individuals hatched in the 1990s. (B) $F_{ROH}$ is correlated with, though much greater than, $F_{ped}$. Dashed line shows 1:1 relationship. (C) Levels of $F_{ROH}$ do not differ between 78 embryos that failed to hatch and 31 hatched individuals born during the same time period. (D) No impact of $F_{ROH}$ on adult survival in a Cox proportional hazards model (p=0.46). See Table S2 for full model output. (E) $F_{ROH}$ is negatively correlated with annual reproductive output, though this relationship is not statistically significant. See Table S3 for full model output. (F) No differences in $F_{ROH}$ between 51 individuals that produced viable offspring at any point in their life versus 14 individuals that never successfully reproduced. Note that horizontal black lines in panels C and F denote the means of each distribution.

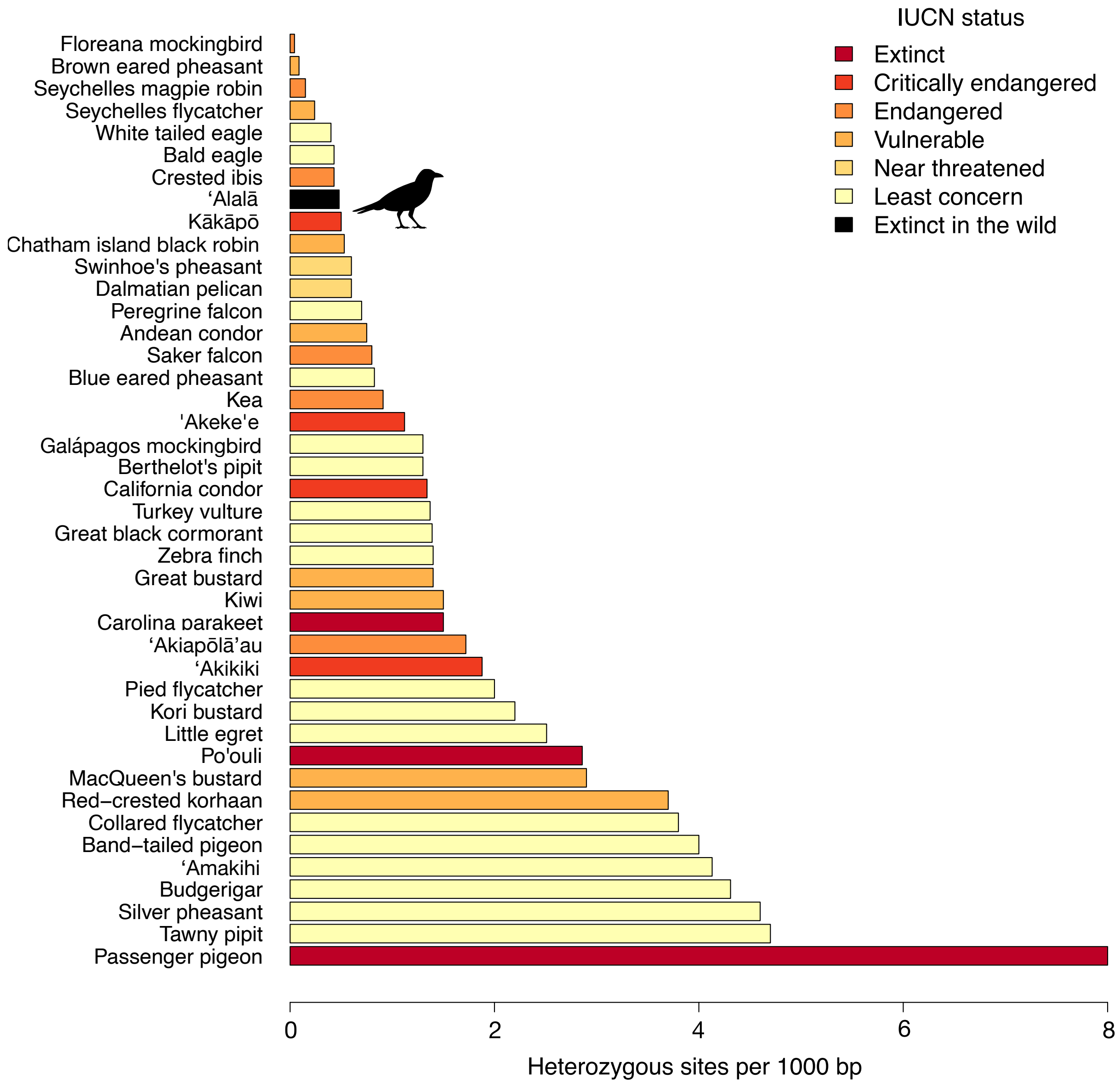


**Figure S5:** Comparison of genome-wide heterozygosity for ‘Alalā to estimates from 42 bird species from previous studies.

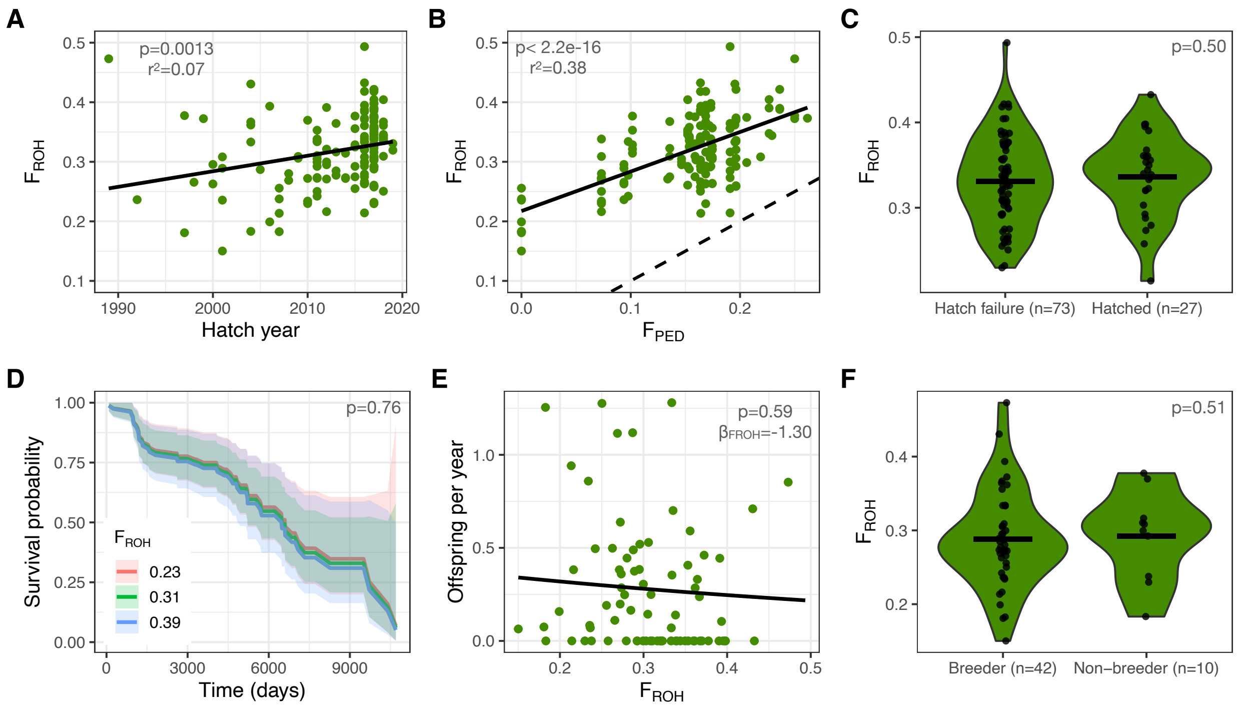


**Figure S6:** Patterns of inbreeding and inbreeding depression in the 'Alalā conservation breeding program when assuming an ROH threshold of >1Mb and removing low coverage (<10x) individuals. (A) Levels of inbreeding (measured by $F_{ROH}$ for ROH > 1Mb) increase over time in hatched individuals (n=81), though are elevated even in the 1990s. (B) $F_{ROH}$ is correlated with, though much greater than, $F_{ped}$. Dashed line represents 1:1 relationship. (C) Levels of $F_{ROH}$ do not differ between 73 embryos that failed to hatch and 27 hatched individuals born during the same time period. (D) No impact of $F_{ROH}$ on adult survival in a Cox proportional hazards model (p=0.76). See Table S2 for full model output. (E) $F_{ROH}$ is negatively correlated with annual reproductive output, though this relationship is not statistically significant. See Table S3 for full model output. (F) No differences in $F_{ROH}$ between 42 individuals that produced viable offspring at any point in their life versus 10 individuals that never successfully reproduced. Note that horizontal black lines in panels C and F denote the means of each distribution.

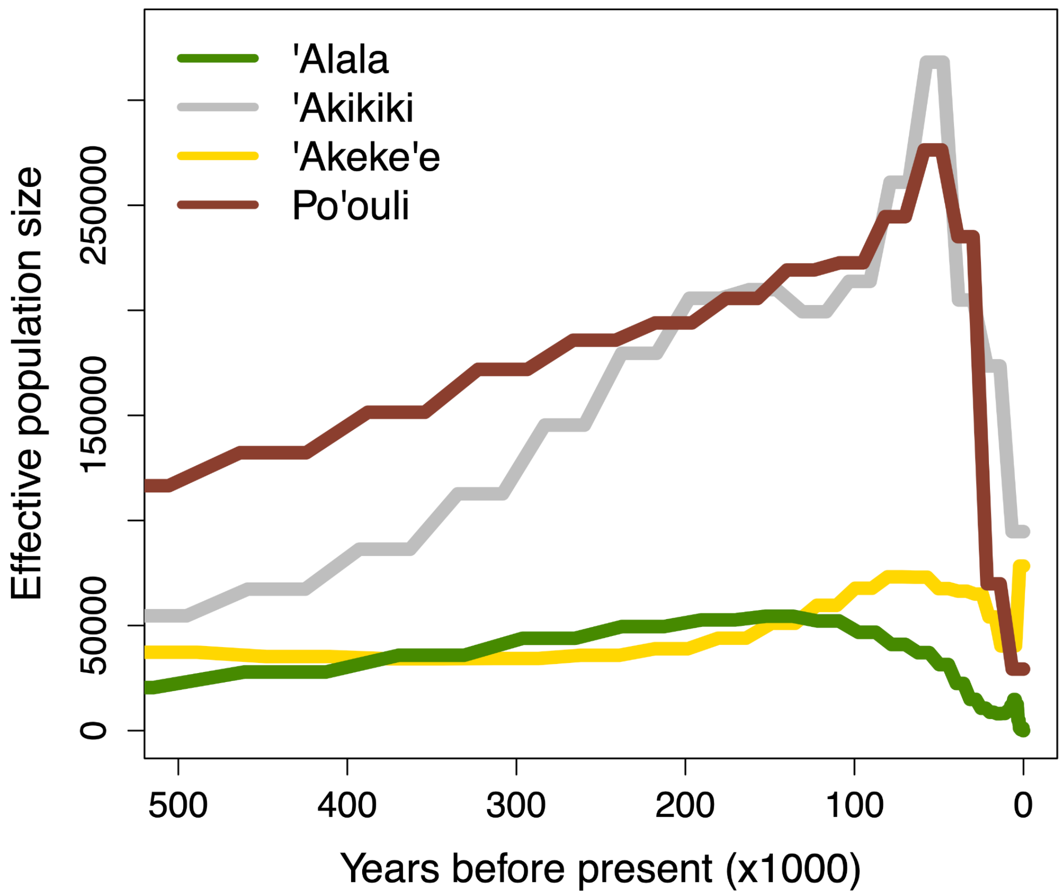


**Figure S7:** Effective population sizes over the last 500,000 years for ʻAlalā in comparison to three Hawaiian honeycreeper species from Kyriazis et al. (2025).

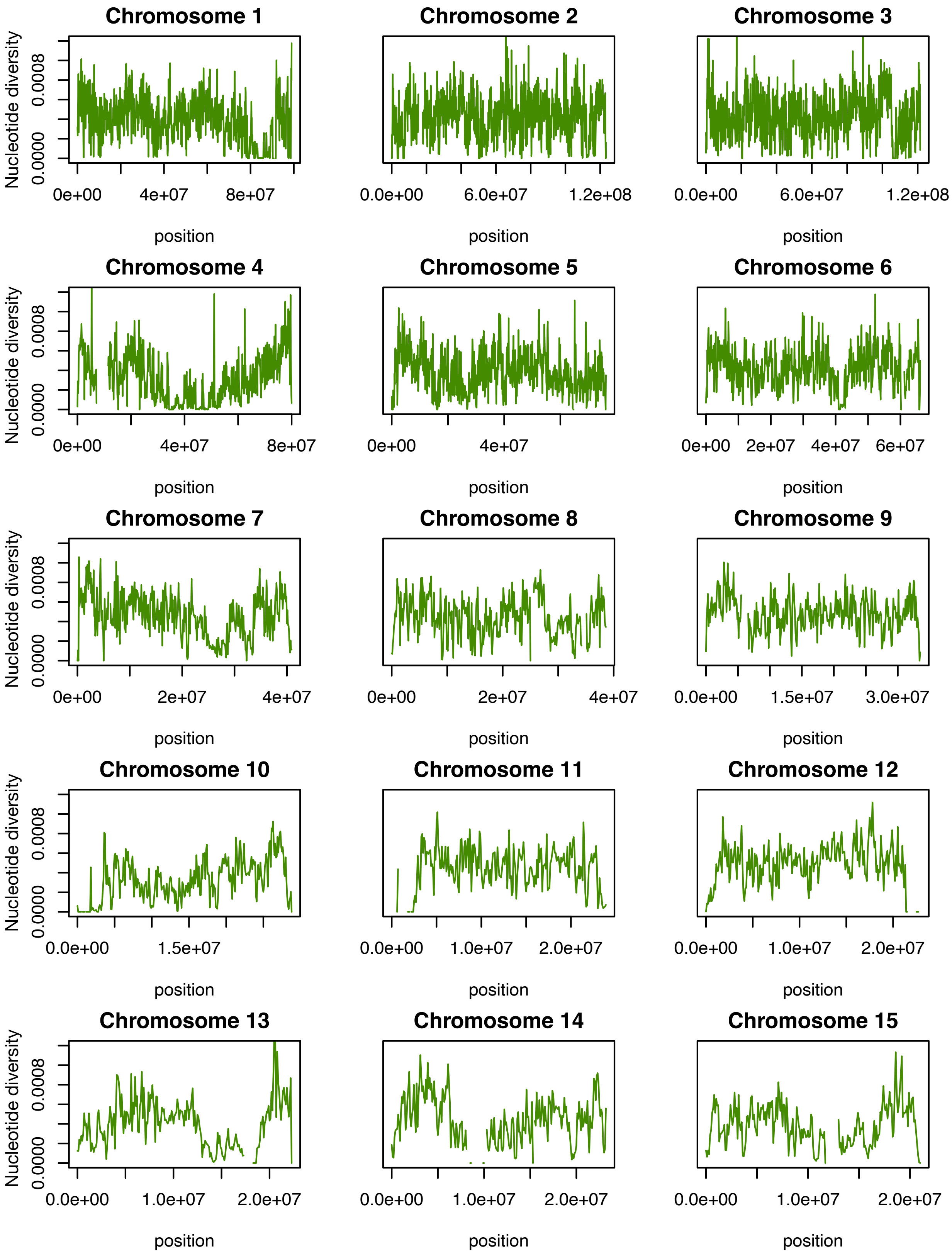


**Figure S8:** Patterns of nucleotide diversity in 100kb windows across the first 15 chromosomes for all living ‘Alalā individuals (n=50).

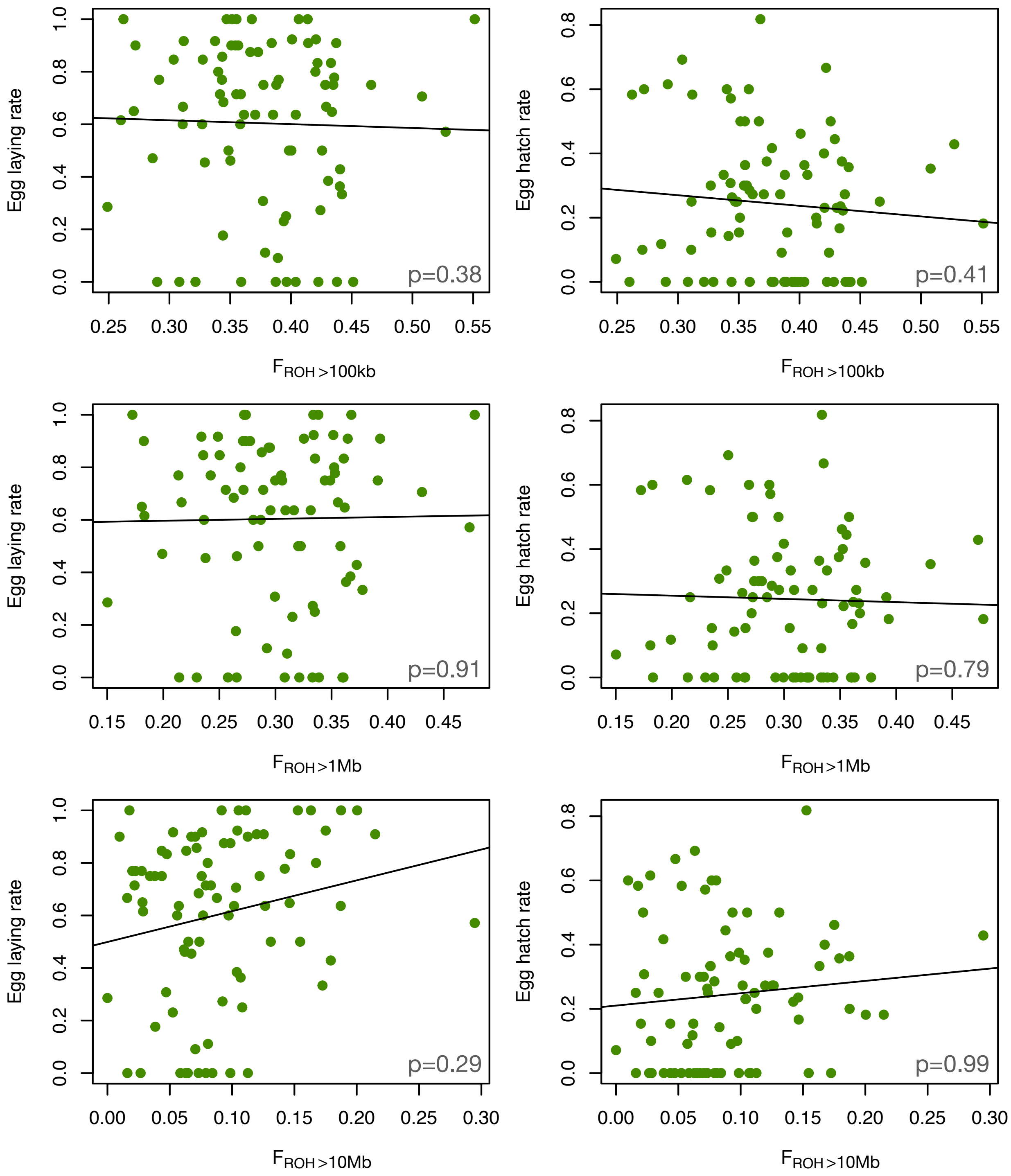


**Figure S9:** No correlation between $F_{ROH}$ and average lifetime egg laying rate or egg hatch rate for 84 individuals that were given opportunities to breed. Top panels show $F_{ROH}$ for ROH > 100kb, middle panels show $F_{ROH}$ for ROH>1Mb, and bottom panels show $F_{ROH}$ for ROH >10 Mb.

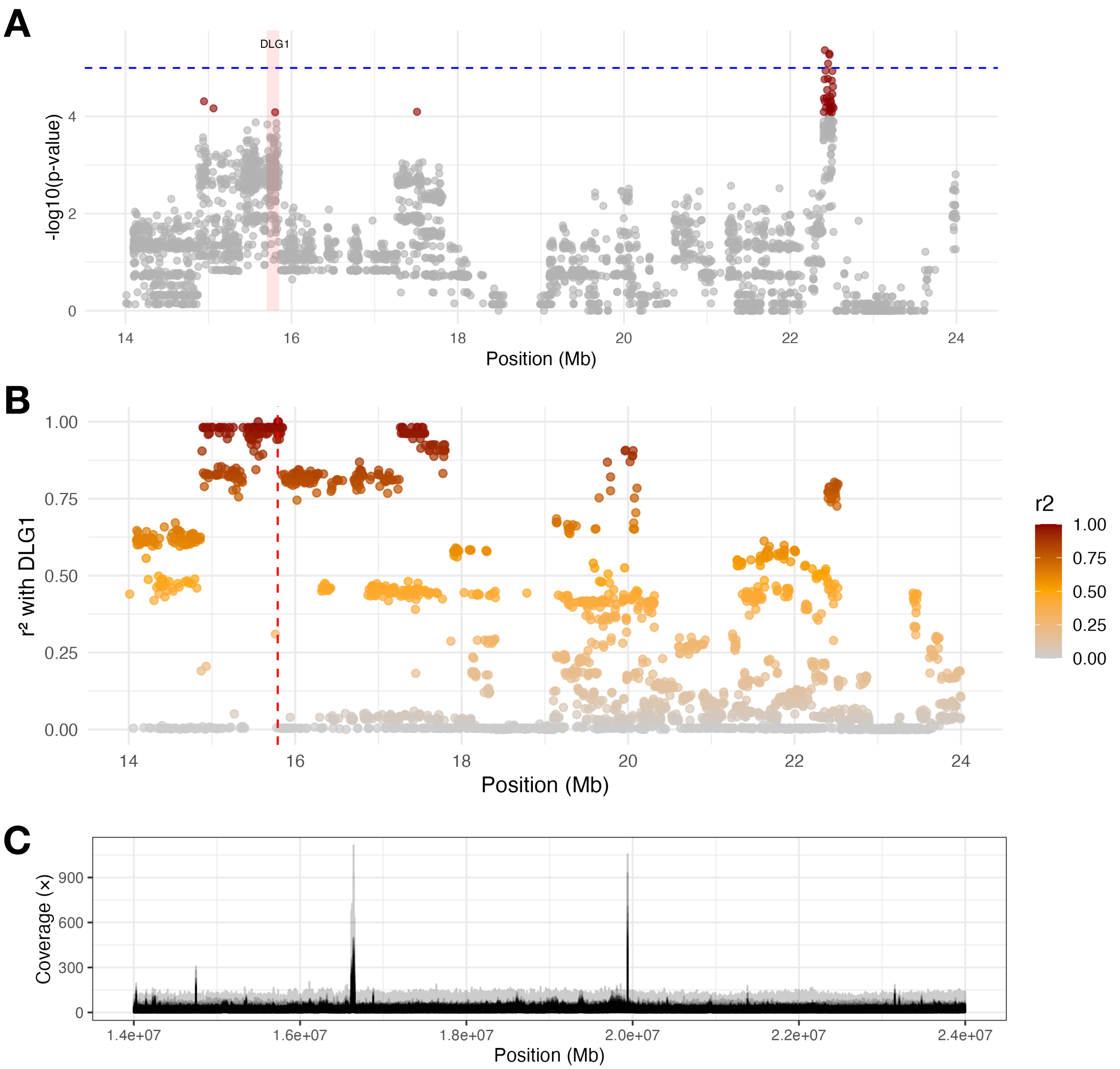


**Figure S10:** Patterns of association, linkage disequilibrium, and coverage near putative recessive lethal allele in DLG1 on chromosome 10. (A) Several suggestive SNPs (p<1e-4) are present on chromosome 10, including in DLG1 (shaded red) and a larger peak within a pseudogene near position 22.5 Mb. (B) Patterns of LD between the putative recessive lethal SNP in DLG1 and the rest of the chromosomal region show a non-monotonic decrease suggestive of structural variation. (C) Depth of coverage plotted for all sequenced individuals show two peaks at ~16.5 Mb and ~20 Mb suggestive of a ~3.5 Mb inversion.

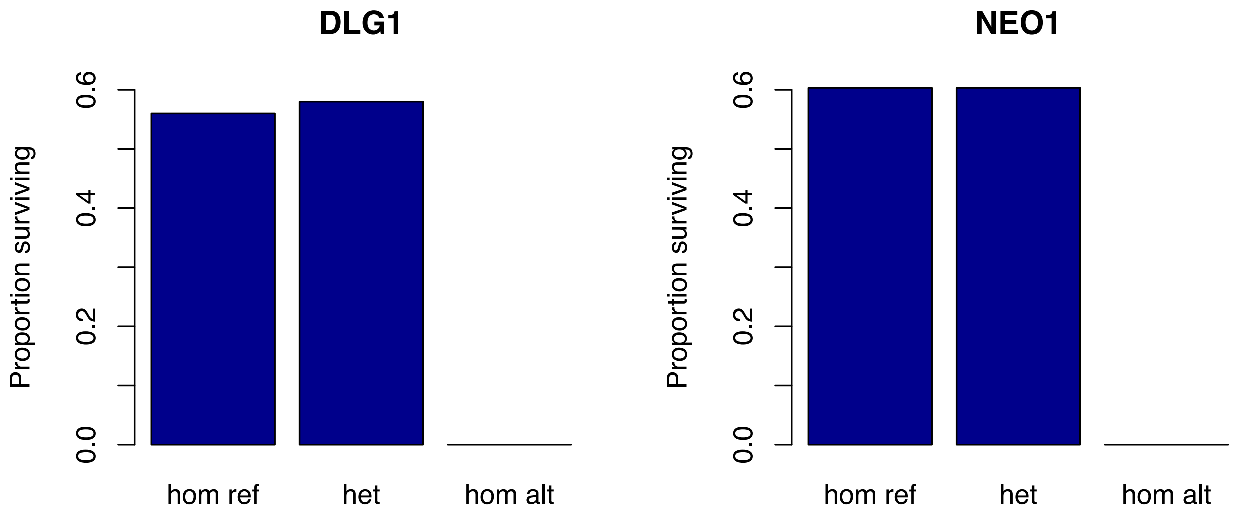


**Figure S11:** Proportion of individuals hatching by genotype for putative recessive lethal variants in DLG1 and NEO1. Note that, in both cases, ~55-60% of homozygous reference and heterozygous individuals hatch whereas 0% of homozygous alternate individuals hatch (0/8 and 0/15, respectively), consistent with expectations for a recessive lethal variant.

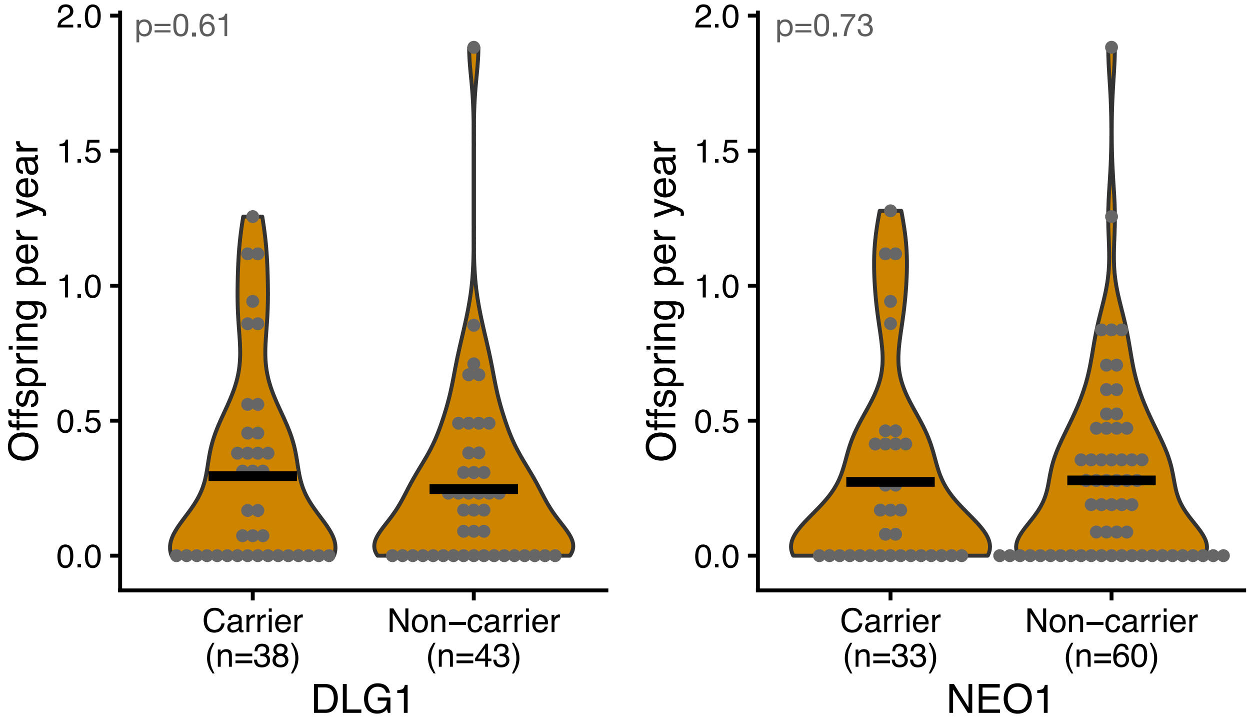


**Figure S12:** No differences in reproductive rates among heterozygous carriers of putative recessive lethal loci and non-carriers for all sequenced individuals. Means shown with horizontal black bar.

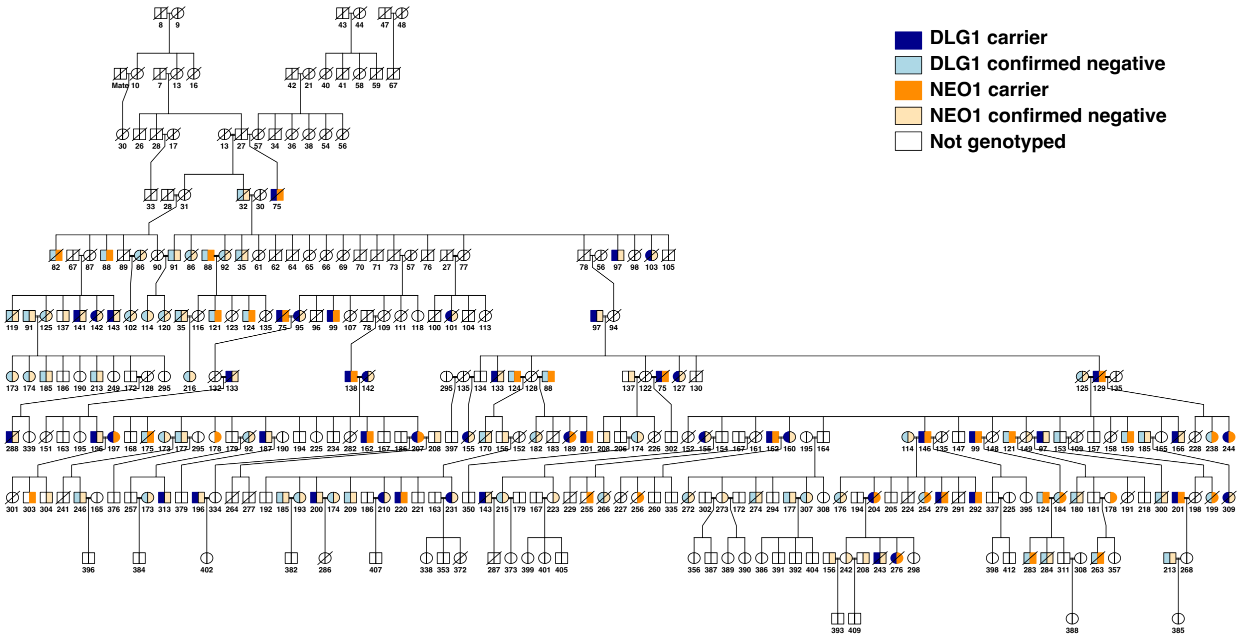


**Figure S13:** Complete pedigree of the captive ʻAlalā population showing recessive lethal genotypes for sequenced individuals. Deceased individuals shown with a black slash.

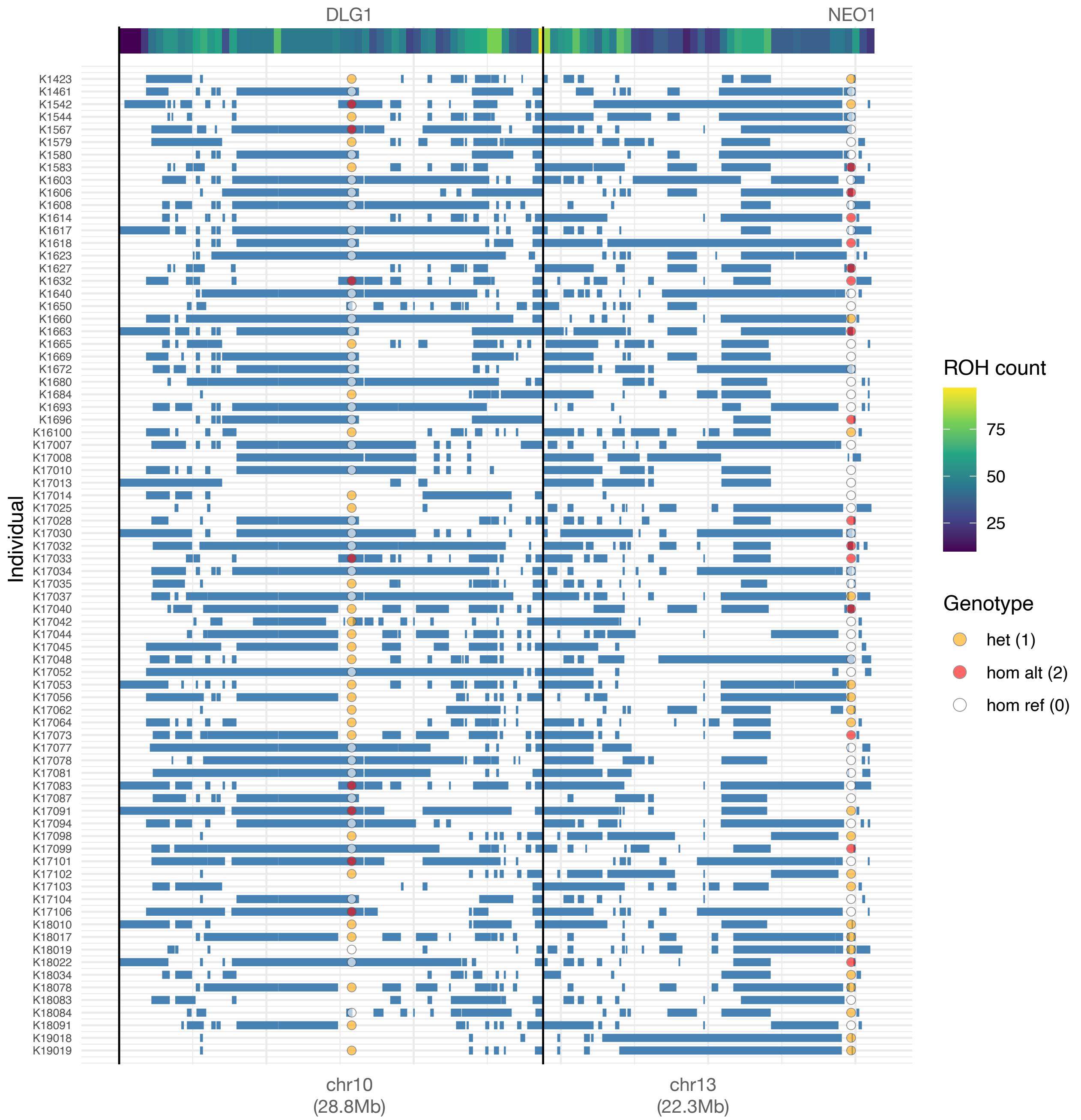


**Figure S14:** Presence of long runs of homozygosity in relationship to putative recessive lethal genotypes on chromosomes 10 and 13. Note that homozygous alternate genotypes are exclusively within long ROH at the DLG1 locus on chromosome 10 but not at the NEO1 locus on chromosome 13.

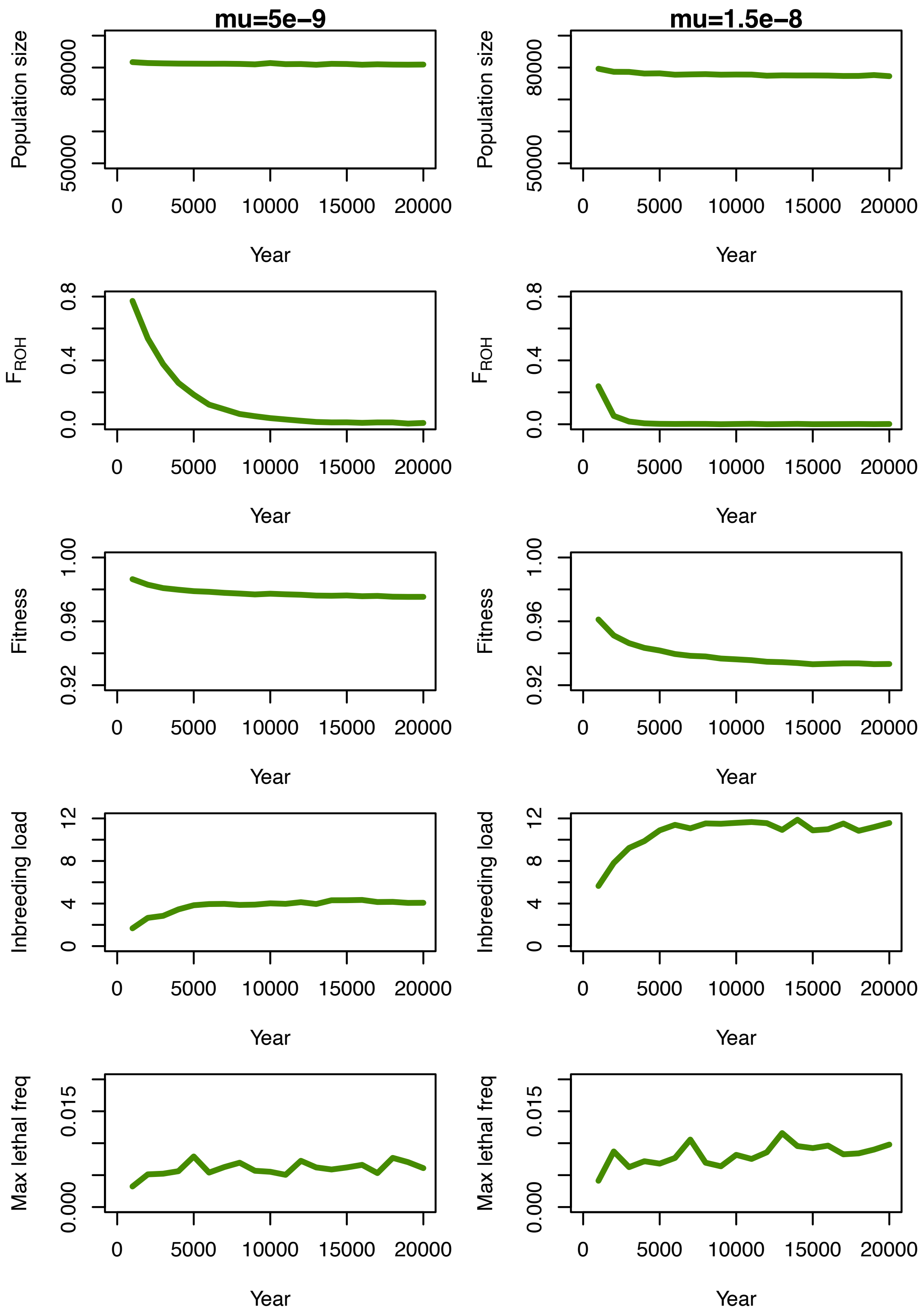


**Figure S15:** Burn-in dynamics for the eco-evolutionary simulation model, including results for a mutation rate of 5e-9 (left panels) and 1.5e-8 (right panels). Top row shows population size during the burn-in, second row shows $F_{ROH}$ for ROH > 1 Mb, third row shows average fitness, fourth row shows average inbreeding load for survival to 6 years, and final row shows maximum recessive lethal allele frequencies.

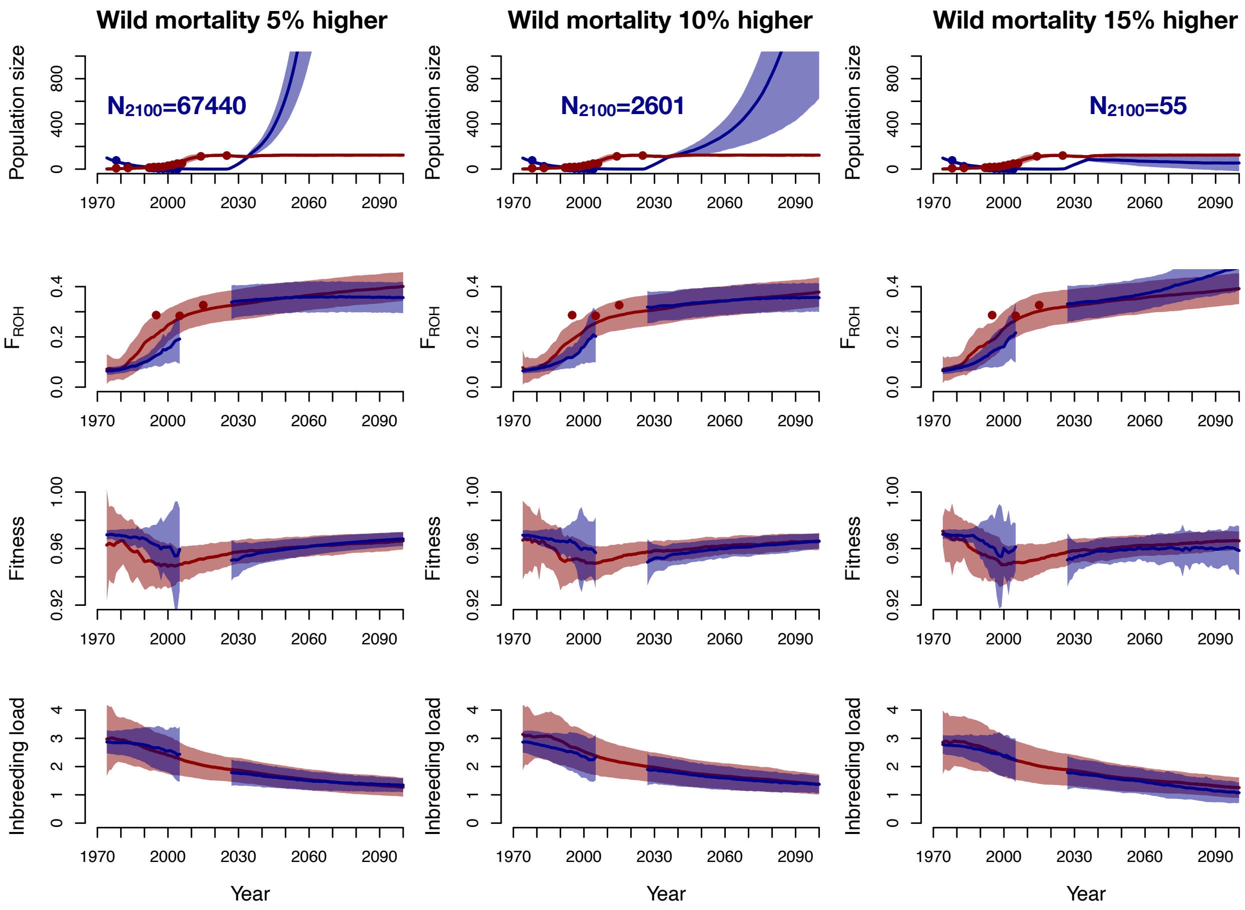


**Figure S16:** Eco-evolutionary simulation results showing impacts of the mortality rate in the wild on recovery. Each column depicts a different relative mortality increase (5%, 10%, 15%) in the wild population (blue) relative to the captive population (red) in a scenario where 100 individuals are released over a ten year period from 2026-2036. For each simulated scenario, the top row shows projected population sizes, the second row shows projected $F_{ROH}$, the third row shows projected fitness, and the bottom row shows projected diploid inbreeding load (2B). Results shown from the founding of the captive population in 1973 projected into the future to 2100, with shading showing one standard deviation.

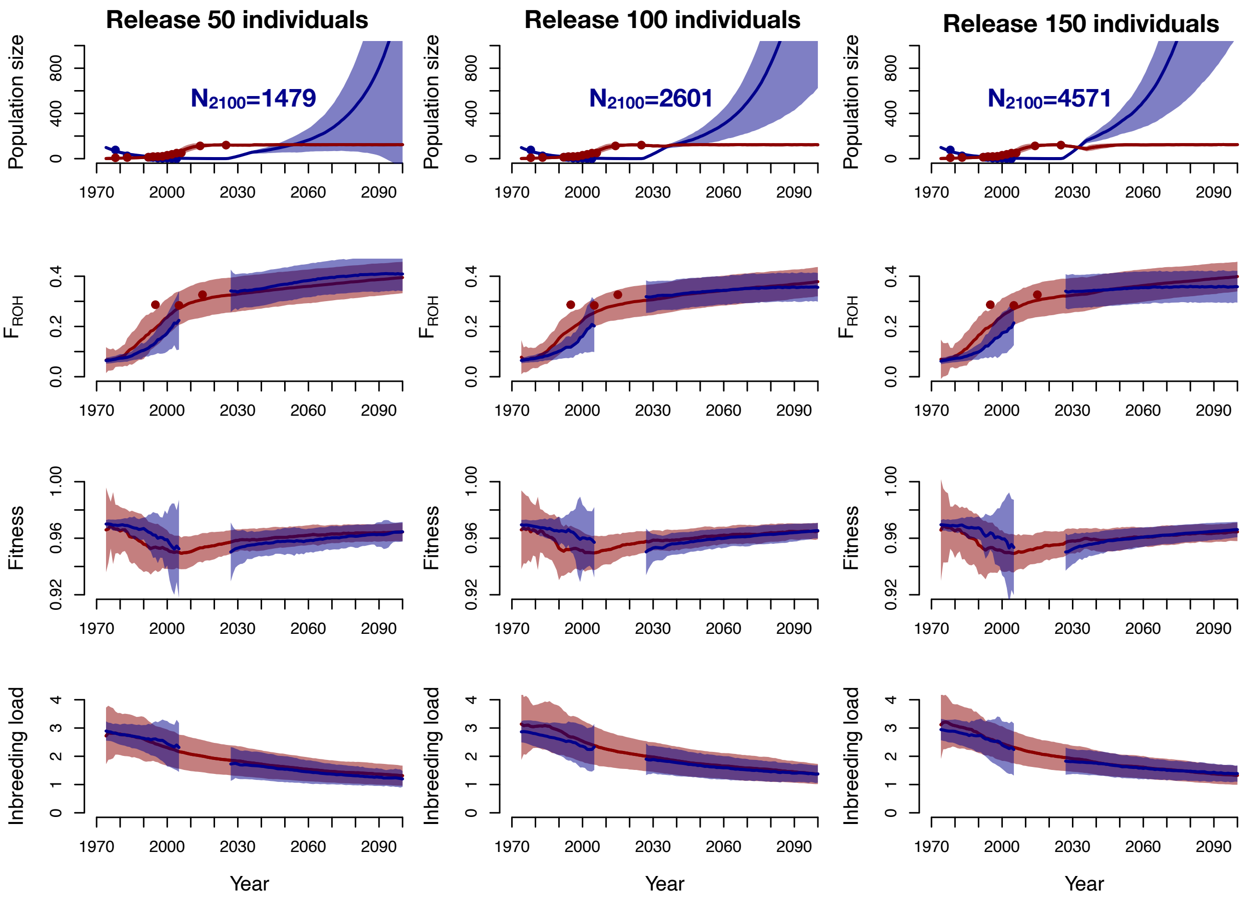


**Figure S17:** Eco-evolutionary simulation results showing impacts of the number of individuals (50, 100, or 150) released over a ten year period from 2026-2036. For all scenarios, we assume an increase in annual mortality rates in the wild population (blue) relative to the captive population (red) of 10%. For each simulated scenario, the top row shows projected population sizes, the second row shows projected $F_{ROH}$, the third row shows projected fitness, and the bottom row shows projected diploid inbreeding load (2B). Results shown from the founding of the captive population in 1973 projected into the future to 2100, with shading showing one standard deviation.

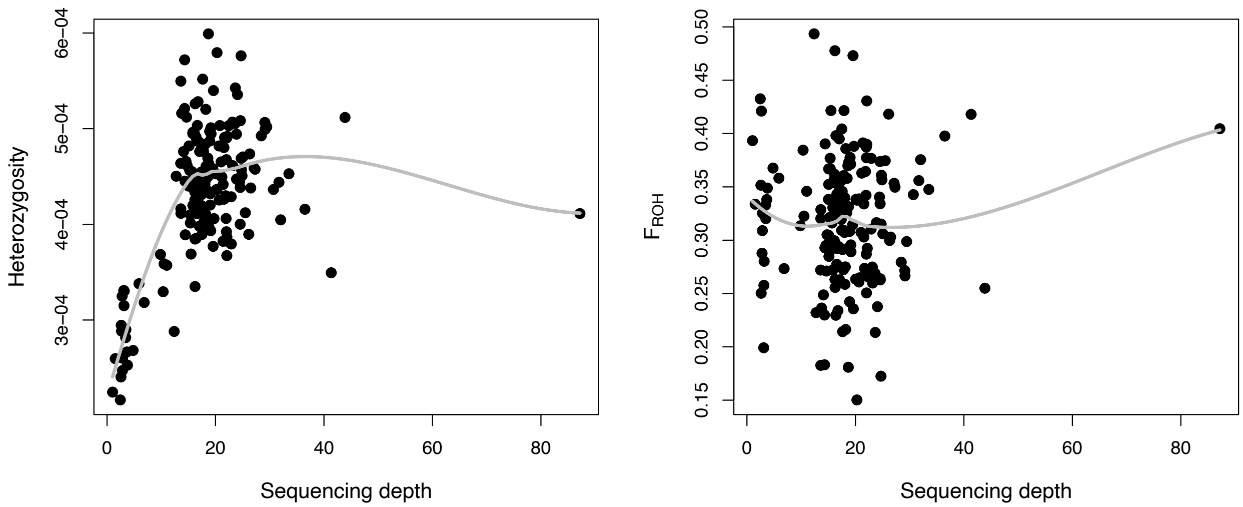

**Figure S18:** Sequencing depth is correlated with individual heterozygosity (left) but not $F_{ROH}$ (right).
